# Ligand Binding Free Energy Landscapes at the Tubulin Colchicine Site from Coarse-Grained Metadynamics

**DOI:** 10.64898/2026.02.24.707696

**Authors:** Andrea Grazzi, Chelsea M. Brown, Maurizio Sironi, Siewert J. Marrink, Stefano Pieraccini

## Abstract

Accessing deeply buried binding sites remains a major challenge in structure-based drug discovery, where accurate description of both protein dynamics and ligand binding pathways is required. Funnel metadynamics enables simulation of complete binding processes but is computationally demanding at the all-atom resolution. By adopting the Martini 3 force field, coarse-grained funnel metadynamics (CG-FMD) substantially reduces computational requirements while retaining enhanced sampling capabilities. In this work, we assess the capability of CG-FMD to model ligand recognition at the deeply buried colchicinoids site of the tubulin ***αβ***-heterodimer, a multisite protein of strategic importance. We investigated the binding of colchicine, podophyllotoxin and combretastatin-A4, recovering free energy profiles with improved statistical convergence compared to AA-FMD and comparable to experimental references. In particular CG-FMD binding free energies present mean absolute errors between 3 and 10 ***kJ mol^−^*^1^**. These results propose CG-FMD as an efficient, physics-based framework for probing ligand binding to challenging sites.

## 1 Introduction

Expanding the scope of druggable targets remains a central objective in the pharmaceutical industry, motivating continuous efforts to identify and exploit deep pockets or cryptic binding sites. From a modeling perspective, binding-site accessibility represents a major challenge. Indeed, many binding sites are cryptic and absent in apo structures, while others are gated by dynamic processes ranging from side-chain rearrangements to large-scale loop motions[1]. Nevertheless, the field has witnessed considerable methodological advances[2, 3]. For instance, Mixed-Solvent molecular dynamics (MSMD) uses a small fraction of an organic cosolvent as a probe[4] for identifying possible interaction sites, although several drawbacks are reported in literature, ranging from phase separation of the two solvents[5] to induced protein unfolding[6]. Another approach for the characterization of challenging binding sites focuses on expanding the conformational ensemble explored by the protein during molecular dynamics (MD) simulations. Meller *et al*.[7] found that extensive All Atom-MD (AA-MD) simulations starting from the crystal apo structure of plasmepsin II did not obtain the desired pocket opening, whereas simulations beginning from a conformational ensemble generated from AlphaFold did capture the phenomenon of interest.

Simulating the complete binding process of a ligand to a poorly accessible site combines the challenges of capturing with sufficient accuracy the chemical interactions involved in the binding pose with sampling the protein’s behavior along the binding pathway. Examples of simulation approaches that model the entire physical pathway have been reviewed by Limongelli[8] and comprise funnel metadynamics (FMD)[9], umbrella sampling[10, 11] and steered molecular dynamics[12]. Metadynamics[13, 14] accelerates the exploration of the phase space by depositing a biasing potential along a selection of Collective Variables (CVs), which are descriptors of the system meant to represent the slow degrees of freedom that hinder exploration of the process of interest (e.g. the distance between the ligand and its binding site). The addition of the biasing potential allows to “fill” the energy minima, allowing diffusive exploration of the free energy landscape. The Well-Tempered (WT) metadynamics implementation[15], upon which FMD is based, reduces the amount of bias progressively added, allowing the simulation to converge more smoothly[16]. FMD introduces a funnel shaped positional restraint, which surrounds the binding site in its conical region, allowing the ligand to freely explore the relevant binding modes. Towards the solvent, the cone is reduced to a narrow cylinder, which still allows the ligand to drift apart from the protein to visit the solvated state, but avoids complete dispersion in the solution bulk and facilitates reapproach to the binding site. This methodology has been applied successfully in literature[17, 18], but it suffers from the drawbacks inherent to AA-MD, namely the computational cost and poor scalability of the system’s size.

Recently, we have shown that coarse-grained funnel metadynamics (CG-FMD) can be an interesting alternative[19]. This approach combines state-of-the-art enhanced sampling capabilities of FMD with the reduced cost of a coarse-grained representation of the system using the Martini 3 model. The Martini 3 force field[20] has proven effective in modelling protein ligand interactions[21] by adopting a mapping scheme that associates groups of atoms to CG-beads with a resolution sufficient to maintain specificity at the level of chemical fragments, as exemplified by several works with soluble proteins[21–23] and transmembrane systems[24–27]. The CG representation of the ligand, where one CG-bead can virtually represent similar fragments, allows to perform MD simulations effectively with a pharmacophoric model of the compound, allowing efficient exploration of the chemical space while still encompassing physics-based contributions from the biosystem, such as protein flexibility and explicit representation of the solvent. With the continuous development of new tools for streamlining the setup of the simulations[28–30], CG-MD is arising as a promising tool for the drug-discovery pipeline[31].

In this work, we tested the ability of the CG-FMD methodology to elucidate the full binding process of three pharmaceutical scaffolds, namely colchicine, podophyllotoxin and combretastatin-A4 (Figure 1a) to the colchicinoids site of the tubulin *αβ*-heterodimer. The tubulin *αβ*-heterodimer is the essential repeating unit in microtubules (MT), which are highly dynamic polymers that regulate a variety of cellular functions. Modulating MT dynamics by targeting tubulin is of strategic interest for the development of therapeutical approaches addressing cancer and neurodegeneration. Indeed, several chemical scaffolds targeting eight topographically distinct binding sites have been experimentally identified[32], denoting the tubulin dimer as a multisite protein. Colchicine[33], podophyllotoxin[33] and combretastatin-A4[34] bind with high affinity to the same deeply buried pocket located at the interface between the two tubulin subunits (Figure 1b), referred to as the colchicinoids site[35].

**Fig. 1.**
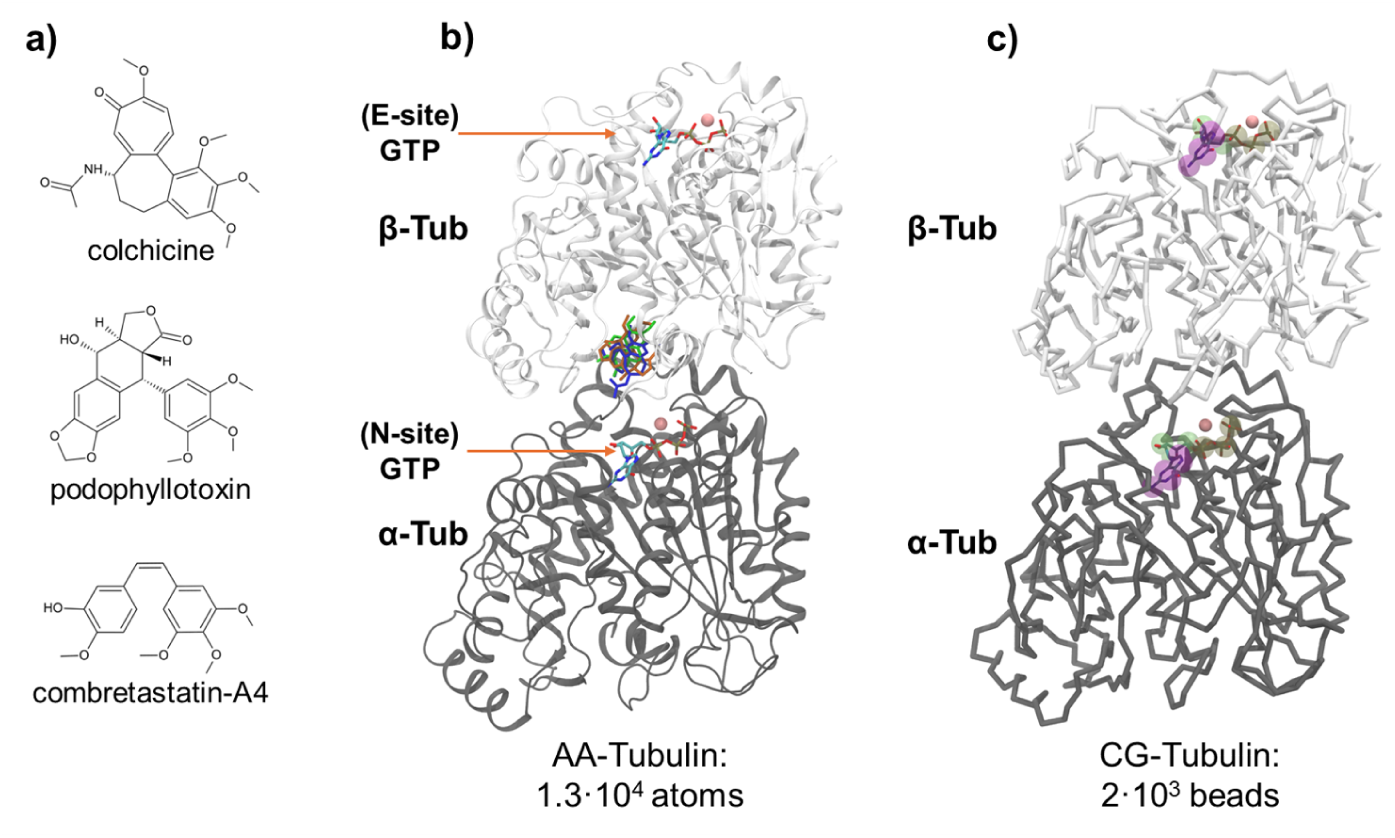
a) Chemical structure and name of the compounds investigated in this work. b) Representation of the tubulin dimer. Crystallographic pose of the investigated ligands is coloured in blue for colchicine, orange for podophyllotoxin and green for combretastatin-A4. c) Representation of the tubulin heterodimer at Martini 3 resolution. For the GTP molecules in the nucleotides sites, an overlay between the CG-model, depicted with transparent spheres, and the reference atomistic pose is reported.

In summary, we show that AA-FMD was able to obtain a free energy landscape of the binding process consistent with experimental evidence, while binding free energy estimation proved difficult to converge. Interestingly, we found that CG-FMD could recover energy landscapes comparable to the atomistic results with a fraction of the computational resource. Furthermore, thanks to the achievable increase in sampling, binding free energies predictions showed better statistical convergence and were comparable to experimental reference with a mean absolute error ranging from 3 to 10 *kJ mol^−^*^1^, depending on the ligand considered. Overall, we propose CG-FMD as a computationally efficient, physics-based approach for the exploration of binding events in deeply buried pockets.

## 2 Results

### 2.1 Equilibrium CG-MD

To evaluate whether known binding sites could be sampled in unbiased simulations, equilibrium MD was performed at Martini 3 resolution. To best reflect solved experimental structures, GTP and magnesium were modelled within their known binding sites. For the target compounds, a small number of molecules were placed in the bulk solvent. Overall, 0.6-0.7 ms of simulation time was collected to provide ample sampling for binding site detection (Figure S17b).

Initial density analyses for the compounds were consistent with the multisite nature of tubulin, showing a plethora of possible interaction sites. Further clustering binding analysis (Figure 2) identified differences between the compounds. For colchicine, we found three poses located at the entrance of the binding channel, with CG-residence times greater than 200 ns (Figure S20b). Similarly, for podophyllotoxin we observed that the first and second most relevant poses are located at the entrance of the path, with a CG-residence time of 500-600 ns (Figure S21b). Finally, for combretastatin-A4, we sampled the full binding process to the crystallographic site, with the top-pose and two others being observed inside the pocket (Figure S22b). A summary of the relation between the top-ten poses determined from unbiased CG-MD across all protein models and the experimentally determined binding site is reported in Figure S19.

**Fig. 2.**
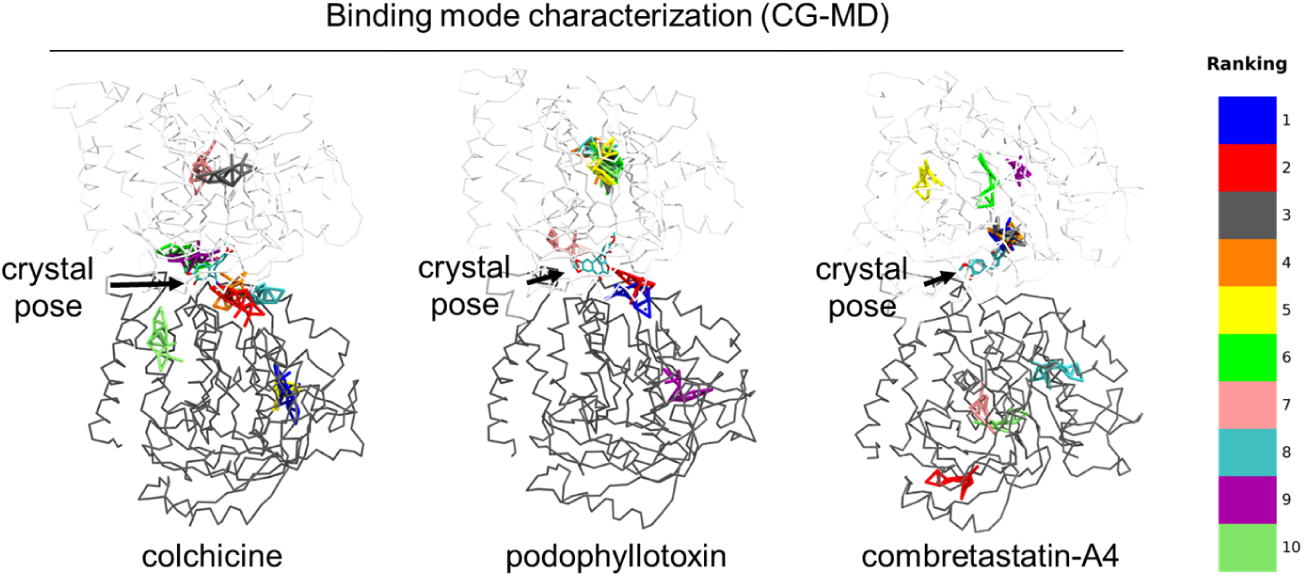
Characterization of tubulin-ligand interaction patterns from equilibrium CG-MD. A) Top-ten representative ligand binding modes (BMs) obtained from simulations with (Olives)-Tubulin are reported in different colours; crystallographic pose is reported in cyan licorice.

Interestingly, we observed that alongside the intrinsic properties of the ligand, such as conformational flexibility and steric hindrance, the CG-network applied to the backbone of the protein impacted results, demonstrating the power of performing equilibrium simulations to understand the flexibility needed in the system to observe binding. Overall equilibrium CG-MD simulations indicated that a primary interaction site for these ligands located at the entrance of the channel for the colchicinoids’ site was accessible, but the full binding event to the crystallographic site proved difficult to observe and to sample extensively with unbiased simulations.

### 2.2 All-Atom Funnel Metadynamics

The first series of simulations to elucidate the complete binding process was conducted with All-Atom Funnel Metadynamics (AA-FMD), starting from an equilibrated pose of the ligand inside the crystallographic binding site. As Collective Variables (CVs), we opted for the distance between the centers of mass (COM) of the ligand and of a selection of residues composing the crystallographic site, and for the angle between two of these residues and the center of mass of the ligand (Figure S23). If the CVs well-approximate the slow degrees of freedom of the system, by applying a bias alongside these CVs, metadynamics can efficiently accelerate the sampling. Figure 3 depicts the funnel shaped restraining potential acting on the COM of the ligand, which allows freedom of exploration inside the binding site, while avoiding ligand dispersion in the solution bulk.

**Fig. 3.**
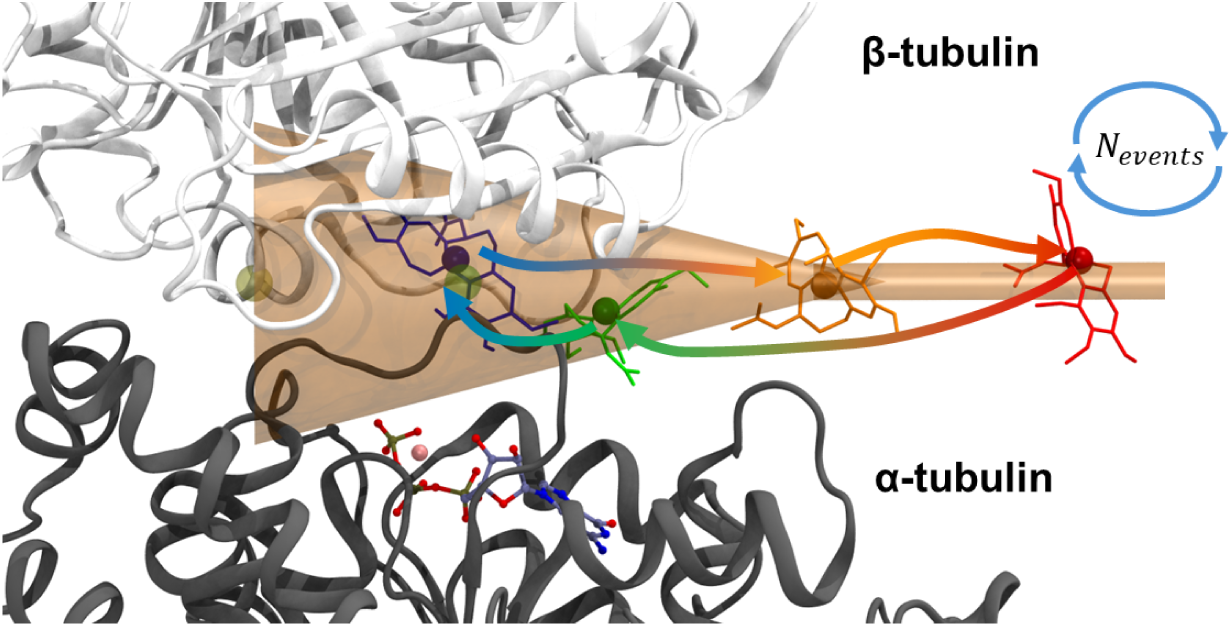
Graphical depiction of the funnel-shaped restraining potential applied to the center of mass of the ligand. For illustrative purposes, several successive snapshots of the ligand colchicine are represented, highlighting how, starting from a putative pose inside the cryptic pocket (depicted in blue), it is possible to observe the full unbinding (poses in orange/red) followed by a reentry in the buried cavity (pose in green).

After optimization of the protocol (see “AA-FMD” in Methods and “FMD-simulation setup” in the Supplementary Information), we obtained converged averaged Free Energy Surfaces (FES) for podophyllotoxin and combretastatin-A4 binding landscapes (first row of Figure 4a), whereas colchicine proved difficult to model. Interestingly, we observed that the topology of the FES is similar across the three ligands considered. In particular, an energy minimum corresponding to the crystallographic pose is correctly identified, and in addition, a secondary basin located at a higher value of ligand-pocket distance (consistent with an entrance pose) is always found. The binding affinity predictions are reported in Table 1 and summarized in Figure 4b. For combretastatin-A4 and podophyllotoxin we obtained estimates that are within 6 *kJ mol^−^*^1^ of the experimental reference, with a standard error of the mean (SEM) of approximately 11 *kJ mol^−^*^1^ across the ten replicas. For colchicine, convergence of the free energy predictions required the extension of the replica duration to 700 ns, yielding a final value within 3 *kJ mol^−^*^1^ from experiments and with a SEM of 12 *kJ mol^−^*^1^

**Fig. 4.**
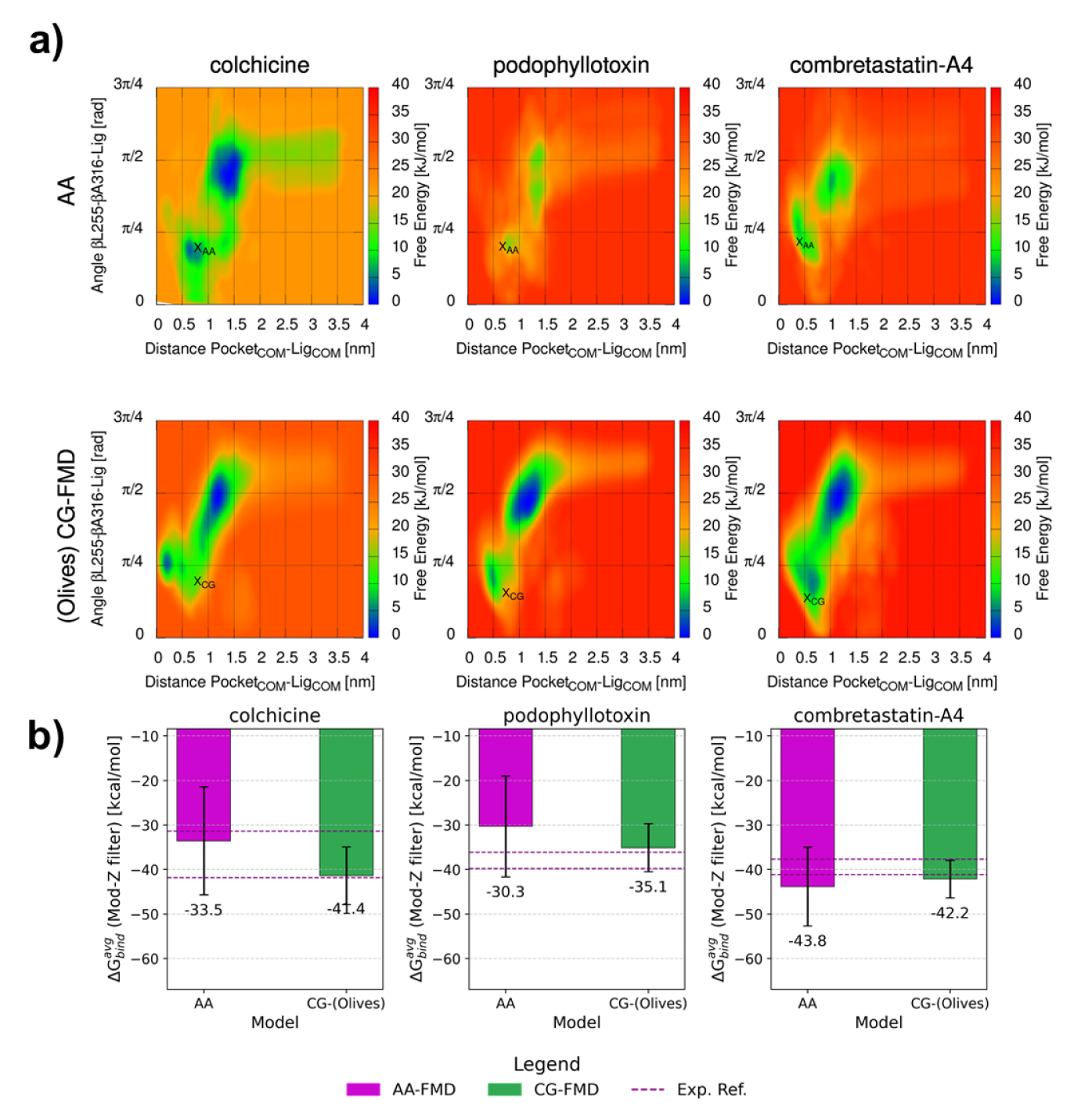
a) Free Energy Surfaces (FESs) obtained from Funnel Metadynamics. Each column corresponds to a different ligand. First row contains results from AA-FMD, while the second row represents data from CG-FMD performed with the Olives CG-networks applied to the backbone of the tubulin dimer. *X_AA_* marks the projection of the crystallographic pose in the CV-space. *X_CG_* indicates the projection of the Martini representation of the crystal pose. b) Summary of the binding free energy estimates obtained for the three ligands investigated in this work. Value reported in purple was obtained with AA-FMD, while results in green were obtained with CG-FMD the Olives CG-network applied to tubulin. Error bars represent the standard error of the mean. Dashed lines indicate the maximum and minimum experimental value reported in literature (summarized in Table S4).

**Table 1.** Summary of binding free energy predictions obtained with Funnel Metadynamics (FMD). For each ligand, results obtained both at the AA and CG resolution are reported, alongside the reference experimental value. Uncertainty over the computational results is the standard error of the mean across the available independent replicas. As preliminary indication of convergence, results obtained at half simulation duration and at full length are reported. CG-results have been obtained with the Olives protein model. Complete results are reported in Table S2, alongside the mean absolute error (MAE) across the different CG-protein networks.

| Ligand | Type | $\Delta G_{\text{bind}}$<br>[kJ mol<sup>-1</sup>]<br>Half-duration | $\Delta\Delta G_{\text{bind}}^{\text{exp-FMD}}$<br>[kJ mol<sup>-1</sup>]<br>Half-duration | $\Delta G_{\text{bind}}$<br>[kJ mol<sup>-1</sup>]<br>Full-duration | $\Delta\Delta G_{\text{bind}}^{\text{exp-FMD}}$<br>[kJ mol<sup>-1</sup>]<br>Full-duration |
| --- | --- | --- | --- | --- | --- |
| Colchicine | Experimental | [-32:-41] | – | [-32:-41] | – |
| | AA-FMD | $-17 \pm 11$ | -19 | $-33 \pm 12$ | -2.8 |
| | CG-FMD (Olives) | $-27 \pm 8$ | -9.2 | $-41 \pm 4$ | 5.1 |
| Podophyllotoxin | Experimental | [-36:-40] | – | [-36:-40] | – |
| | AA-FMD | $-32 \pm 11$ | -5.3 | $-30 \pm 11$ | -6.5 |
| | CG-FMD (Olives) | $-41 \pm 10$ | 3.8 | $-35 \pm 5$ | -2.6 |
| Combretastatin-A4 | Experimental | [-38:-41] | – | [-38:-41] | – |
| | AA-FMD | $-72 \pm 20$ | 32.8 | $-44 \pm 9$ | 4.2 |
| | CG-FMD (Olives) | $-45 \pm 5$ | 5.8 | $-42 \pm 4$ | 2.6 |

Overall, AA-FMD predictions show good agreement with experimental data, exhibiting deviations comparable to those observed in recent studies[17–19, 36](summarized in Table S3).

### 2.3 Coarse Grained Funnel Metadynamics

Results from the AA-FMD simulations suggested that the modelling of the complete binding process to the buried colchicinoids site was possible, albeit at the cost of significant hardware requirements (Table 2) and with challenges in the convergence of the binding free energy estimates. To assess whether comparable results could be achieved with a more computationally efficient approach, we set out to optimize a simulation protocol based on CG-FMD. Concerning the choice of CVs, we decided to maintain the same CVs applied in the AA-FMD campaign, to maintain consistency and ensure comparability of the computational results.

**Table 2.** Summary of the computational performances obtained for AA-FMD and CG-FMD. The last column reports the estimated physical wall-clock time required to complete a simulation of minimal duration that yields acceptably converged results. Hypothesized simulation durations, inferred from examples in this work, are reported in italics.

| Simulation type | CPU-allocation | GPU | Performance [ns/day] | Minimal Wall time [duration] |
| --- | --- | --- | --- | --- |
| CG-FMD | 16-cores<sup>1</sup> | none | 1800 | 66h [*5 μs*] |
| AA-FMD | 8-cores<sup>2</sup> | 1<sup>3</sup> | 80 | 150h [*500 ns*] |

<sup>1</sup>AMD EPYC 7763@2.45 GHz
<sup>2</sup>Intel Xeon Platinum 8358@2.60GHz
<sup>3</sup>NVIDIA Ampere A100 64GB

Interestingly, we found that the FESs obtained with CG-FMD are topologically similar to the predictions of AA-FMD, as reported in Figure 4b for the Olives CG-Network and in Figure S27 for the complete dataset. The morphology of the free energy basins is somewhat shallower, consistent with the notion well established in literature that the potential energy surface underlying a coarse-grained representation of a system is smoother compared to an atomistic one[37]. By applying the biasing potential, CG-FMD simulations were able to sample extensively both the basin located at the entrance of the binding channel and the interior of the colchicinoids site.

Concerning the binding free energy estimates, results obtained with Olives-tubulin are reported in Table 1. Interestingly, we observed that CG-FMD predictions are in good agreement with the experimental reference, yielding a difference between 3 and 5 *kJ mol^−^*^1^. If we consider the results achieved with the other CG-networks (reported in Figure S28 and Table S2), two considerations arise. Firstly, switching from a rigid elastic network to a more flexible protein model consistently improved the estimates, highlighting once more the importance of capturing a more realistic protein behaviour in simulations. Secondly, while there are some differences among the CG-estimates, the overall results do not vary significantly. Indeed, the Mean Absolute Error (MAE) of the CG-FMD estimates ranges from 2.5 *kJ mol^−^*^1^ for combretastatin to 9.5 *kJ mol^−^*^1^ for podophyllotoxin. This suggests that the simulation protocol is solid and can yield reproducible results with some degree of tolerance of the simulation settings.

### 2.4 Exploration of the colchicinoids pocket with CG-MD

The simulation campaign with CG-FMD highlighted that the modelling of the full binding process to the cryptic colchicinoids site was possible, allowing to explore, in addition to the entrance of the channel already frequently visited from unbiased simulations starting from solution bulk (Figure 2), the more buried part of the pocket. Since the resulting FESs are obtained from the combination of two significant perturbations from equilibrium MD, namely the application of Well-Tempered metadynamics, and the introduction of a funnel-shaped restraining potential, we set out to assess whether the free energy basins identified with CG-FMD corresponded to actual energy minima (within the Martini 3 force field framework) or whether they were simulation artefacts. To this end, we performed several plain CG-MD simulations (Table S5) starting from poses already inside the pocket. Distance and angle CVs have been computed with the same definition used for FMD and their evolution in time has been projected over the FES obtained from CG-FMD, as reported for a subset of simulations with the Olives CG-model in Figure 5 (full dataset represented in Figures S32-34).

**Fig. 5.**
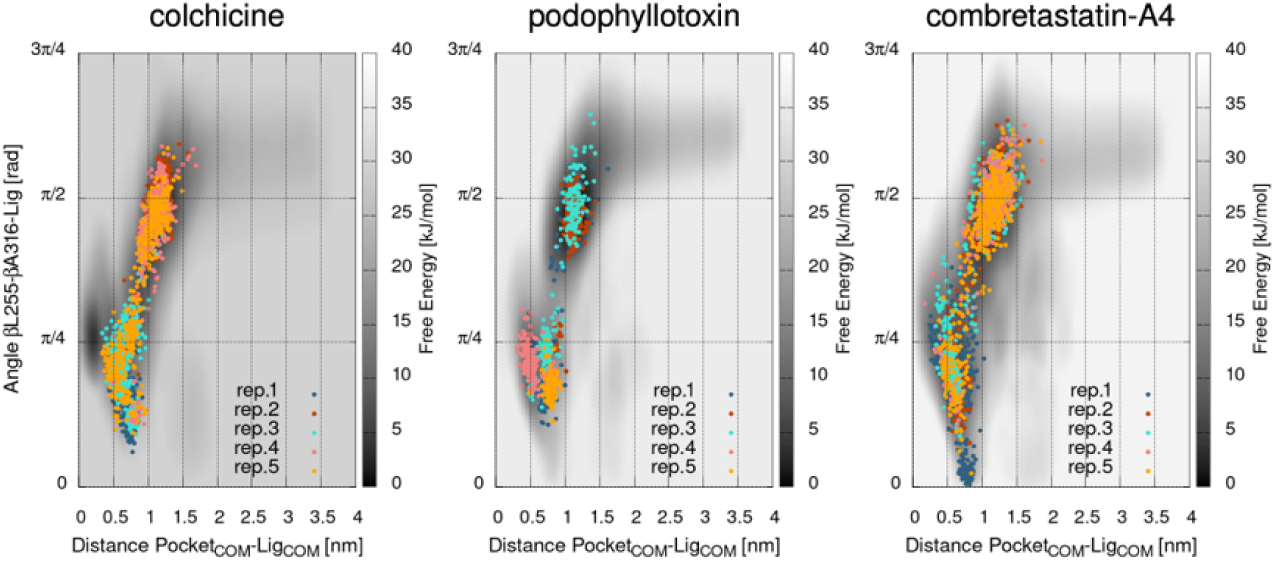
Validation of CG-FMD FES through equilibrium CG-MD starting from a ligand pose inside the binding site. Each plot represents the overlap between equilibrium CV-timeseries (colored dotted series) and the FES obtained from CG-FMD (grayscale). These results have been obtained with the Olives CG-network applied to the tubulin’s backbone, Results for the other CG-models are reported in Supplementary Information, “Equilibrium CG-MD exploration of the pocket”

Remarkably, we observed that equilibrium simulations visited the energy basins predicted by CG-FMD, providing an internal validation for the prediction obtained from the enhanced sampling protocol.

## 3 Discussion and conclusions

The objective of this work was to assess the feasibility of modelling the binding process of small molecules of pharmaceutical interest to the deep colchicinoids site of the tubulin *αβ*-heterodimer. Unbiased CG-MD simulations were typically able to identify several long-lived, ligand binding modes located at the entrance of the experimentally known binding site, but the full penetration was seldom observed and not for all the chemical scaffolds considered. To elucidate the full phenomenon, we decided to apply funnel metadynamics as enhanced sampling methodology. Atomistic results presented a contrasting picture. On the one hand, after optimization of the simulation protocol, we were able to obtain a final average FES identifying the entrance basin and the crystallographic basin as global minima. On the other hand, estimation of the binding free energy highlighted that convergence across the dataset of ten simulations for each ligand was challenging to achieve. In particular, the minimum duration for achieving convergence in two out of three cases was approximately 300 ns (as illustrated in Figure S30), with the third instance (colchicine) requiring the extension of the simulations from 500 ns to 700 ns to reach an estimate compatible with experimental references.

In contrast, convergence of the statistical estimate of CG-FMD was more readily achieved, as reported in Figure S31. For instance, affinity predictions for combretastatin-A4 seem to converge after 4 *µ*s. For colchicine and podophyllotoxin, the initial estimates tend to oscillate for the first few microseconds, while they stabilize from 5 *µ*s. Indeed, an estimation of the binding affinity for these two ligands at half simulation duration, is not too dissimilar from the more consolidated value after the entire 10 *µ*s (as reported in Table 1), which can be a useful insight for the investigator looking for preliminary results without committing to long simulation times. Indeed, the MAE observed after just 5 *µ*s of CG-FMD, ranging from 5 to 10 *kJ mol^−^*^1^ (Table S2) depending on the ligand, should be compared with the spread of 4 to 10 *kJ mol^−^*^1^ in the values of experimental binding affinities reported for the three ligands (summarized in Table S4).

In closing, we would like to emphasize the advantages in terms of computational resources offered by CG-FMD compared to AA-FMD. Similarly to our previous work[19], we estimated a binding event acquisition rate *λ* by fitting the average number of observed binding events against the wall time necessary to perform the simulation (Figure S36). Interestingly, CG simulations are fifteen to thirty times faster than atomistic simulations in collecting binding events (Table 3). This implies that for a given amount of wall time, CG-FMD can estimate ligand binding free energies based on a dataset of events that is potentially larger by at least one order of magnitude compared to AA-FMD. Consequently, the significant reduction in statistical uncertainty can partially compensate for the loss of accuracy that derives from the CG-representation of the biosystem. Furthermore, CG-FMD presents significantly lower hardware requirements (Table 2), enabling greater scalability of targets. An interesting point of comparison is offered by the work of Shan *et al.*[38]. They set out to investigate the binding modes of four compounds to interleukin-2 by performing unbiased AA-MD. After performing an extensive simulation campaign of 134 *µ*s on the Anton HPC infrastructure, they observed multiple binding modes on the surface of the protein besides the crystallographic one, resorting to free energy perturbation (FEP) calculations to assess whether the forcefield could correctly rank the native pose. Interestingly they found that while the pose most similar to the crystal had often the lowest energy, the binding affinity FEP predictions (ranging from −20 to −10 *kcal mol^−^*^1^) were significantly different from experimental data (ranging from −11 to −5 *kcal mol^−^*^1^). Considering that the system size for the tubulin dimer is at least ten times larger (2.5 *·* 10^5^ atoms against the 2.3 *·* 10^4^ atoms for the system of Shan *et al* [38]), replicating that protocol at the AA resolution would require a substantial commitment of computational resources, with the challenge of obtaining converged results. Indeed, Gaspari *et al.*[34] reported an uncertainty of 2 *kcal mol^−^*^1^ in the free energy prediction related to the investigation of combretastatin-A4 unbinding from tubulin with WT-metadynamics. On the other hand, simulating at Martini 3 resolution allowed us to obtain free energy landscapes and binding free energy predictions well correlated to the experimental evidence at a fraction of the cost.

**Table 3.** Comparison of the binding event acquisition rate (expressed in wall time), between AA-FMD and CG-FMD. Results obtained with *t_cooldown_* = 2 *ns* and *t < min_duration_* = 4 *ns* in the binding events counting protocol. Additional details on the statistical fit are provided in Figure S35

| Ligand | AA-FMD<br>$\lambda$ [Events/h] | CG-FMD (EN)<br>$\lambda$ [Events/h] | CG-FMD (Olives)<br>$\lambda$ [Events/h] | CG-FMD (goMartini)<br>$\lambda$ [Events/h] |
| --- | --- | --- | --- | --- |
| colchicine | 0.03±0.01 | 0.63±0.01 | 0.66±0.01 | 0.76±0.01 |
| podophyllotoxin | 0.02±0.01 | 0.64±0.01 | 0.60±0.01 | 0.68±0.01 |
| combretastatin-A4 | 0.04±0.01 | 0.88±0.01 | 1.31±0.03 | 1.44±0.03 |

Overall, we propose that CG-FMD can be applied as an efficient simulation approach for preliminary investigations of deep binding sites, combining the reduced computational cost of modelling a complex biosystem at Martini 3 resolution with the advantages of a state-of-the-art enhanced sampling approach, such as FMD.

## 4 Methods

### 4.1 Ligand parameterization

Martini 3 parameters for the nucleotide GTP and the ligands podophyllotoxin and combretastatin-A4 have been defined, whereas parameters for colchicine were taken from a previous work[19]. The parametrization protocol followed guidelines established in literature[20, 39]. Modelling of structural and conformational properties of the small molecules requires the definition of bonded terms, such as bond, angles and dihedral angles. To this end, a reference atomistic simulation is collected and mapped to CG-resolution by associating to each group of atoms one bead with the Center of Geometry approach[20, 39]. Chemical properties of the CG-ligand are defined by the attribution of types to the beads and validated by computing water-octanol partition coefficients. Details on the reference atomistic simulation, as well as validation of the Martini 3 parameters are reported in the Supplementary Information (“Ligand parametrization”).

### 4.2 Protein modelling

Modelling protein behavior is an important aspect of CG-MD simulations since a set of additional bonded terms is required to maintain elements of secondary and tertiary structure[28, 40]. To assess the role of the CG-protein model on the computational findings, we generated parameters for the three established models, namely Elastic Network (EN[41], based on elastic restraints derived from a reference structure), Olives[42] and goMartini[43] (based on Lennard-Jones potential). The validation protocol of these CG-models, which required the collection of reference AA-MD simulations of tubulin, followed literature guidelines[42, 43] and is detailed in the Supplementary Information (“Protein Modelling”). We note on passing that the goMartini model used in the simulation campaign was reparametrized to improve fidelity to AA-MD reference.

### 4.3 Nucleotide modelling

The physiological state of the tubulin *αβ*-heterodimer is characterized by the presence of GTP in both nucleotide binding sites. For the N-site in particular, the accurate modelling of GTP behavior is crucial since this site is located quite close to the colchicinoids’ pocket that we aimed to investigate. In order to avoid excessive nucleotide displacement from its experimental location during the microseconds-long CG simulation, a series of residue-GTP distance restraints was derived from AA-MD and applied to the CG-model of the N-site. In particular, the available AA-MD trajectories of the tubulin dimer have been mapped to CG-resolution (Figure S13a,b) and a contact analysis has been performed between each putative CG-bead of the mapped representation of GTP and the surrounding residues. An average occupancy per residue has been defined as the fraction of simulation frames in which the distance between the ligand bead and the residue fell below the standard cutoff of 0.6 nm implemented in GROMACS. This protocol has been applied to both nucleotide sites, allowing us to quantify the most important interactions needed to maintain a GTP pose consistent with crystallographic evidence (Figure S13c). A subset of long-lived interactions has been selected for the definition of the harmonic distance restraints to be applied during CG-MD simulations, and their average distance has been taken as the reference value.

### 4.4 Equilibrium CG-MD from solution bulk

The CG-tubulin dimer was solvated with regular water beads and NaCl has been added to neutralize the systems’ charge and reach a physiological concentration of 0.15 M. Four to six ligand molecules were randomly placed in the solution bulk. Final systems’ size and composition are reported in Figure S17a. After energy minimization with the steepest descent algorithm, a first NPT equilibration with the Berendsen barostat[44] (*τ_P_* =4.0 ps, compressibility 3 *·* 10*^−^*^4^ *bar^−^*^1^), the v-rescale thermostat[45] (*τ_T_* =1.0 ps) and position restraints on the backbone beads was performed. Then 30-35 replicas (Figure S17b) have been performed under NPT ensemble (Parrinello-Rahaman barostat[46], *τ_P_* =12.0 ps, compressibility 3 *·* 10*^−^*^4^ *bar^−^*^1^) with randomization of the initial velocities and a duration of 20.05 *µ*s (initial 50ns have been considered as equilibration and discarded from subsequent analysis). Neighbor-list settings have been modified according to the recommendations of Kim et al[47]. In particular, we set verlet-buffer-tolerance=-1, rlist=1.35, nstlist=nsttcouple=nstpcouple=20. VMD[48] was used to compute ligand occupancies with the Volmap tool. To elucidate the interaction patterns and characterize the relevant binding modes, a custom GROMACS pipeline was developed. Similarly to the approach by Bartocci *et al* [24], the protocol first detects long-lived events by performing a protein-ligand contact analysis. Then, a cluster analysis is applied to identify representative binding modes (Figure S14). Further details are reported in the Supplementary Information (“Characterization of ligand binding modes”).

### 4.5 AA-FMD

The starting conformation for AA-FMD simulations was taken from holo simulations after 10 ns of equilibration under NPT conditions starting from the crystallographic ligand pose (PDB ID 4O2B for colchicine[49], 1SA1 for podophyllotoxin[33], 5LYJ for combretastatin-A4[34]). The FMAP-GUI was used for the definition of the funnel shaped potential[50]. A distance CV was defined between the COM of the ligand and the COM of a group of residues lining the interior of the colchicinoids pocket. An angle CV was constructed among the COM of the residues *β*-Leu255, *β*-Ala316 and of the ligand, as described in Figure S23. Well-Tempered metadynamics was applied to the distance-angle CV pair with bias deposition rate every 500 *md_steps_* (1 ps at 2 fs), initial hills height of 3.0 *kJ mol^−^*^1^, biasfactor=24 and simulation duration of 500 ns. To avoid protein denaturation upon ligand extraction, an upper restraint of 0.3 nm was imposed on the RMSD computed over tubulin’s backbone. Furthermore, after observing that metadynamics strain could negatively affect the stability loops involved in the CVs definition (Figure S25a,b), a second set of RMSD restraint of 0.2 nm was imposed on local portions of the binding site. Considering the two residues involved in the angle CV definition (*β*-Leu255, *β*-Ala316), a sequence interval of *i ±* 3 was chosen to identify the portion of the loops to be stabilized (Figure S25c).

After reweighting, a free energy surface (FES) was computed for each replica. The estimated ligand binding constant was computed with a correction for the presence of the funnel-shaped potential[50]:

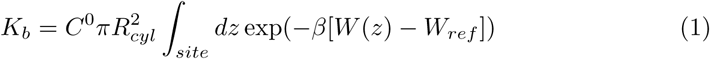

Here, *C*^0^ is the standard concentration equal to 1/1.661 *Å*^−3^, *R* is the radius of the cylinder towards the solvent, *W* (*z*) is the value of the FES inside the basin of integration, *W_ref_* is the free energy value in the unbound state and *β* is the inverse of the product between the simulated temperature T and the Boltzmann constant *k_B_*. For the definition of the basin boundaries, the projection of the ligand crystallographic pose in the CV-space was considered ( Figure S26). The binding constant was converted to a free energy value using 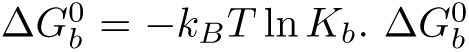 values from independent replicas have been averaged, yielding a final binding affinity estimate with an associated statistical uncertainty defined as the standard error of the mean. When computing the average, an outlier removal criterion based on the modified Z-score was applied.

### 4.6 CG-FMD

To maintain consistency and comparability with the atomistic results, the simulation protocol for CG-FMD was based on the same funnel geometry and CVs definition. The bias deposition rate was kept every 500 *md_steps_*(10 ps at 20 fs), and the initial hills height at 3.0 *kJ mol^−^*^1^. Since the binding to the deep pocket could involve the crossing of energy barriers, we set out to optimize the protocol by testing several values of the biasfactor parameter, which regulates the rate of decrease of the metadynamics bias. In particular, we considered three setups: fast-decreasing (biasfactor=12), intermediate (biasfactor=24) and slowly decreasing (biasfactor=50). Concerning the stability of the CG-protein and of the loops defining the CVs, an upper restraint of 0.38 nm was imposed on the RMSD of the entire protein backbone for the Olives and goMartini simulation setups. This was based on the results obtained during the validation of the CG-protein models, which suggested that an RMSD plateau of 0.35-0.38 nm was physiological to (Olives)-tubulin (Figure S9) and (goMartini)-tubulin (Figure S11), and therefore acceptable also during metadynamics simulations. On the contrary, (EN)-tubulin was inherently more stable and no RMSD restraint was needed. No additional loops restraints were imposed, since the CG-protein models already stabilized to an acceptable degree local secondary structure elements. For each combination of CG-protein model, ligand and metadynamics biasfactor, several independent replicas have been collected with a simulation duration of 10 *µ*s (Table S1).

### 4.7 Equilibrium CG-MD inside the colchicinoids’ site

The atomistic structure file obtained from the conversion of the experimental crystal structure of colchicine, podophyllotoxin and combretastatin-A4 was mapped to CG-resolution. 23000 water beads have been added to solvate the tubulin dimer in addition to 0.15 M of NaCl. An energy minimization and a first NPT equilibration of 10 ns with the Berendsen barostat and with position restraints on both the ligand and the backbone beads has been performed to allow the solvent to settle. A second NPT equilibration of 50 ns was performed without restraints on the ligand. Finally, a production simulation of either 10 *µ*s or 20 *µ*s with the Parrinello Rahman barostat was collected (Table S5). This protocol has been repeated independently ten times for each ligand. In order to avoid ligand dispersion in the solution bulk upon hypothetical exit from the binding site, the same funnel restraint introduced in CG-FMD was applied. We note on passing that the conical section of this funnel is large enough to encompass the entirety of the colchicinoids site, so it did not affect the exploration of the pocket at equilibrium.

### 4.8 Assessment of simulation efficiency

The simulation efficiency of CG-FMD compared to AA-FMD has been assessed by computing a binding events acquisition rate *λ*. The ligand was annotated as bound whenever its CVs fell within the intervals denoting the colchicinoids binding site (Figure S26). To accommodate small oscillations around the boundaries, a smoothing procedure (reported previously[19]) has been applied. The number of binding events observed for a certain simulation setup has been averaged across the available independent replicas. The average value *η*, computed at intervals of 50 ns, has been fitted both against the simulated time (*t_md_* = *n_md−steps_· dt_md_*) and against the physical wall time using linear regression. Results from the fitting procedure are reported in Figure S35 and Figure S36.

### 4.9 HPC environment

AA-FMD simulations have been performed using GROMACS[51] 2022.3 patched with PLUMED v2.8.1[52, 53] on LEONARDO Booster nodes (NVIDIA A100)[54]. CG-FMD and equilibrium CG-MD simulations have been carried out using GROMACS v2023.3 patched with PLUMED v2.9.0 on CPU-nodes of Hábŕok cluster, hosted by the Center for Information Technology of the University of Groningen. All other simulations have been performed using GROMACS v2023.5 on desk workstations.

## 5 Data availability

Martini 3 parameters for the protein and ligand investigated in this work, as well as the starting simulation box for eq-CG-MD, AA-FMD and CG-FMD, are provided in a compressed folder (ZIP), available at [55].

## 6 Code availability

The custom code developed for the characterization of the ligand binding modes from equilibrium simulation is available at [56].

## Supplementary Information

Supplementary Information: validation of ligand parameterization, additional data on proteins’ modelling, details on equilibium CG-MD setup, additional data on AA-FMD and CG-FMD, assessment of simulation efficiency (PDF)

## Acknowledgements

We thank the Center for Information Technology of the University of Groningen for their support and for providing access to the Hábŕok high performance computing cluster. We acknowledge ISCRA for awarding this project access to the LEONARDO supercomputer, owned by the EuroHPC Joint Undertaking, hosted by CINECA (Italy). This research was supported by the University of Milan (PSR 2023 Linea 2).

## Author contributions

A.G. performed the simulations and analyzed the data.

A.G. and C.M.B. wrote the original draft. M.S., S.J.M. and S.P. supervised the project and revised the manuscript.

