## supplementary material for "Ligand Binding Free Energy Landscapes at the Tubulin Colchicine Site from Coarse-Grained Metadynamics"

**4 Equilibrium CG-MD exploration of the pocket** **39**

**5 Assessment of simulation efficiency** **43**

### 1 Supplementary Methods

#### 1.1 Ligand Parameterization

##### 1.1.1 Bonded parameters definition

Starting atomistic conformation for the compounds to be parameterized has been retrieved from the RCSB databank. Parameters for the CHARMM36-FF have been generated with the CGENFF python script[1, 2]. Each compound has been solvated with 4500 TIP3P water molecules and in the case of GTP, sodium cations have been included to neutralize the system's charge. After energy minimization with the steepest descent algorithm and a first NPT equilibration for 10 ns at 300 K (V-rescale thermostat[3], $\tau_T$=0.1 ps) and 1 bar pressure (Berendsen barostat[4], $\tau_P$=5.0 ps, compressibility of $4.5 \cdot 10^{-5}$ $bar^{-1}$), production simulations of 1.050 $\mu$s (Parrinello-Rahman barostat[5], $\tau_P$=5.0 ps, compressibility of $4.5 \cdot 10^{-5}$ $bar^{-1}$) have been collected. These trajectories have been converted to CG-resolution using an AA to CG mapping and the center of geometry approach[6, 7]. Reference probability distributions for the bonded terms have been generated from the mapped trajectories. CG-bonded terms have been defined to maximize the overlap with the atomistic reference, as reported in Figure S2,S3,S4. An assessment of the global structural properties has been performed by computing the relative average error on the distribution of the solvent accessible surface area (Figure S5) and by inspecting the Connolly surface (Figure S6). We found good agreement in the reproduction of the molecular shape and the relative error on the SASA was well-below the 5% threshold commonly accepted in literature.

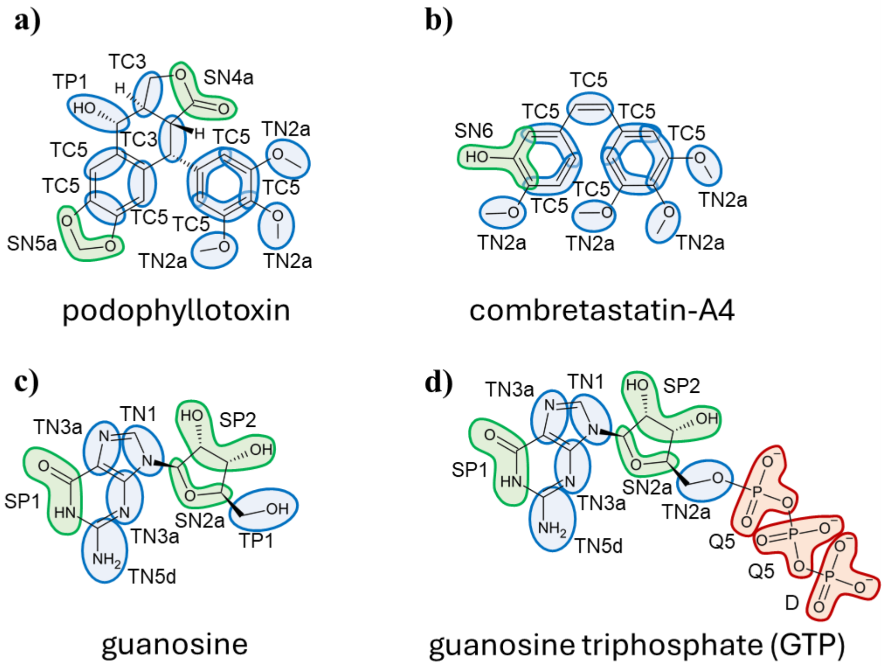

**Supplementary Figure S1** Ligand mapping and bead types attribution scheme for the newly parameterized compounds.

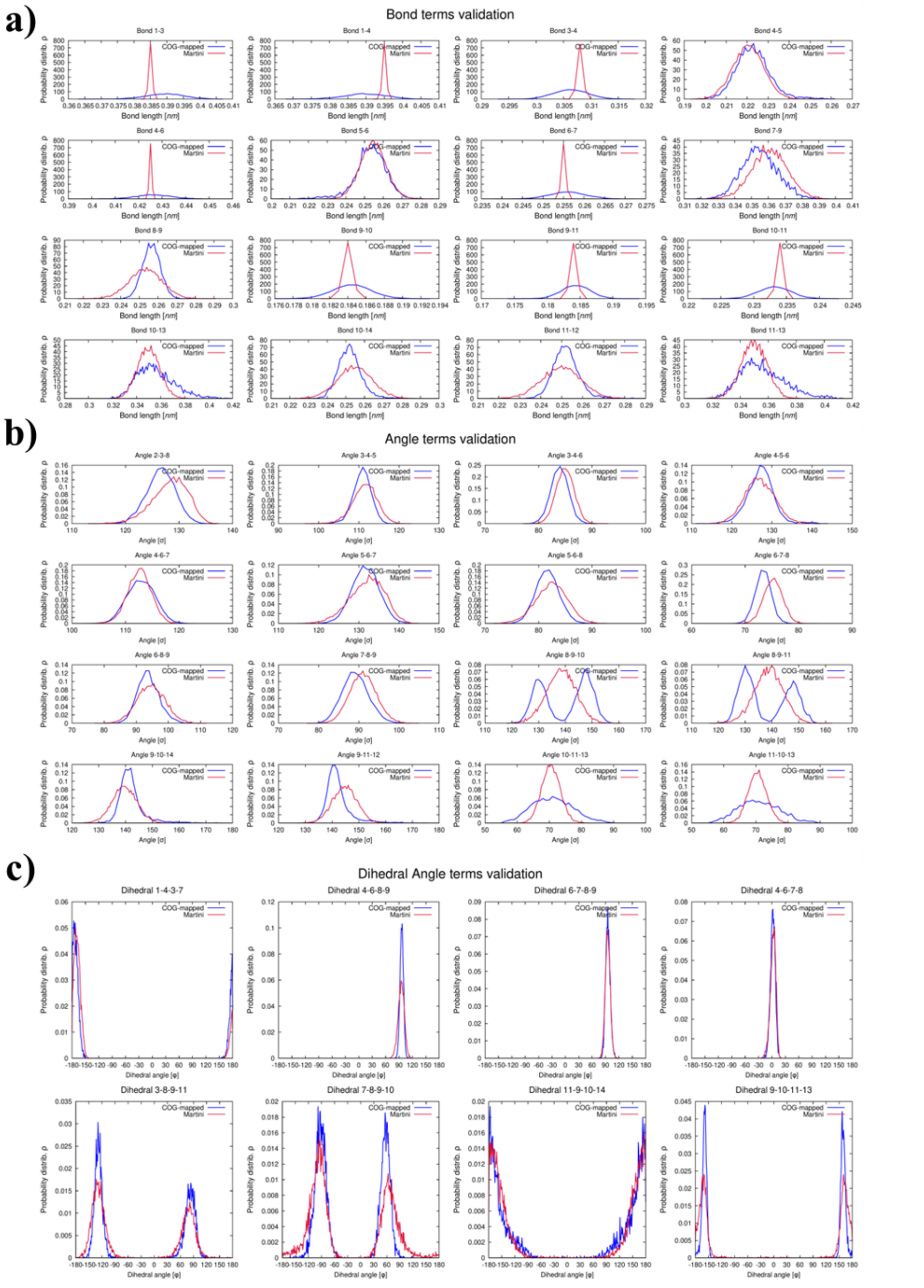

**Supplementary Figure S2** Validation of bonded terms for the CG-model of podophyllotoxin: a) bond terms; b) angle terms; c) dihedral angle terms. Reference probability distribution from atomistic trajectory mapped to CG is depicted in blue. Red distribution has been obtained with the proposed parameterizations of the Martini 3 model of the molecule.

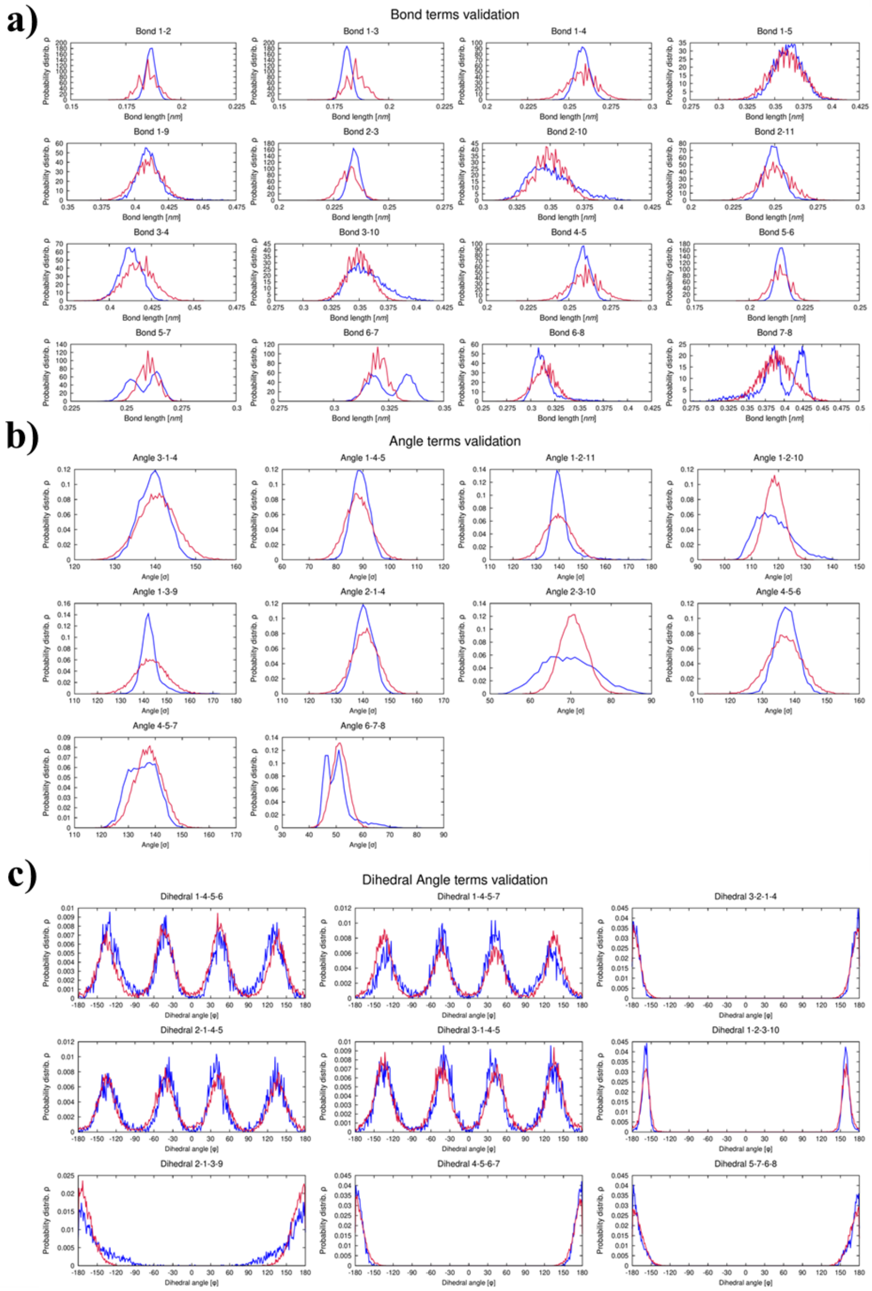

**Supplementary Figure S3** Validation of bonded terms for the CG-model of combretastatin-A4: a) bond terms; b) angle terms; c) dihedral angle terms. Reference probability distribution from atomistic trajectory mapped to CG is depicted in blue. Red distribution has been obtained with the proposed parameterizations of the Martini 3 model of the molecule.

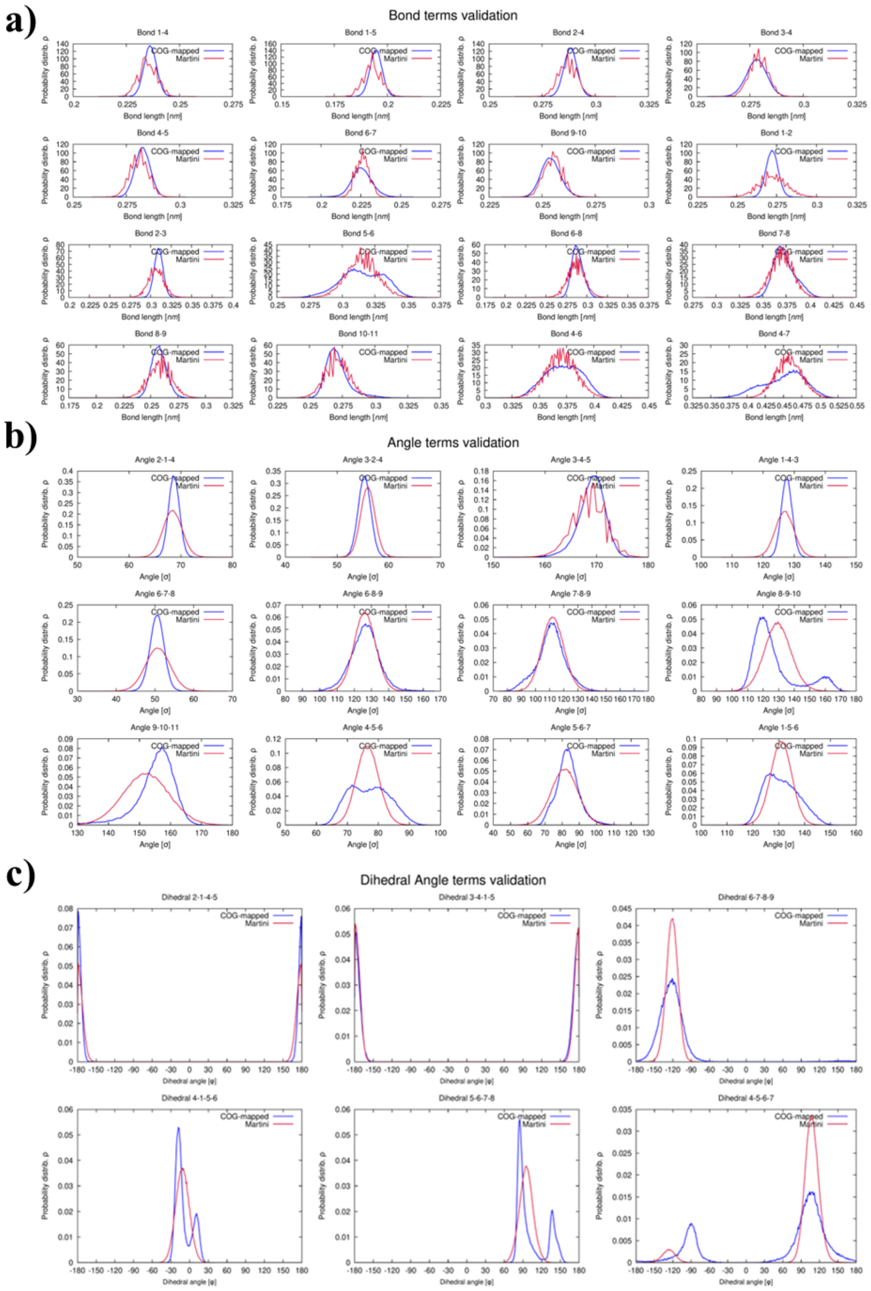

**Supplementary Figure S4** Validation of bonded terms for the CG-model of GTP: a) bond terms; b) angle terms; c) dihedral angle terms. Reference probability distribution from atomistic trajectory mapped to CG is depicted in blue. Red distribution has been obtained with the proposed parameterizations of the Martini 3 model of the molecule.

**a)**

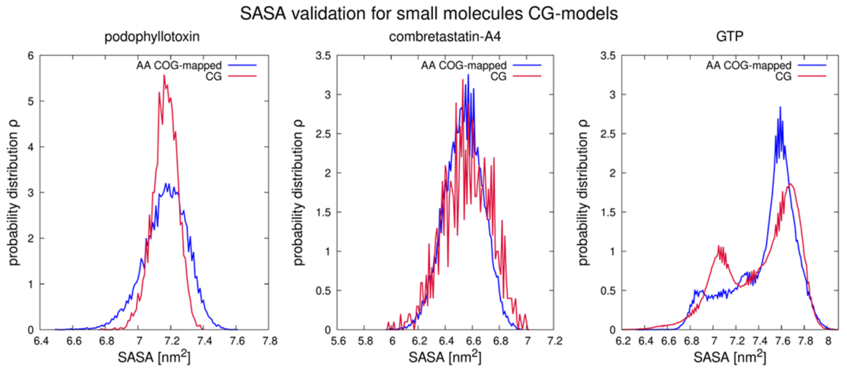

**b)**

| SASA validation | | | |
|---|---|---|---|
| *Compound* | AA-simulation [nm$^2$] | CG-simulation [nm$^2$] | Err% |
| podophyllotoxin | 7.2±0.1 | 7.2±0.1 | 0.1% |
| combretastatin-A4 | 6.5±0.1 | 6.6±0.2 | 0.4% |
| GTP | 7.5±0.3 | 7.4±0.3 | -0.8% |

**Supplementary Figure S5** Validation of the Solvent Accessible Surface Area (SASA). a) Graphical representation of the SASA distribution from a reference AA trajectory (blue) and from a simulation of the CG-model of the parameterized small molecule (red). b) Average values of the two distributions are reported with their respective relative errors on the difference.

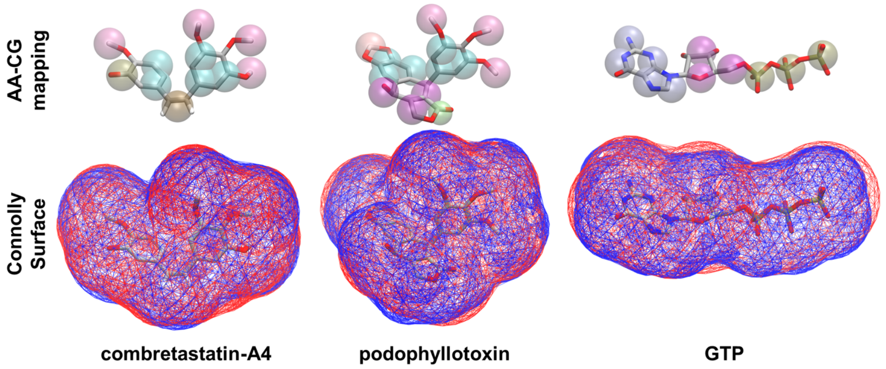

**Supplementary Figure S6** Validation of molecular shape. Top row: overlap between an atomistic conformation energy-minimized (licorice) and the corresponding Martini 3 representation (transparent spheres). Bottom row: overlap between the Connolly surface computed over the atomistic structure (blue) and the surface calculated over the coarse-grained model (red).

##### 1.1.2 Non-bonded parameters definition

The chemical properties of the CG-model of a compound are determined by the types attributed to its beads (Figure S1). The bead-type attribution scheme is typically validated by computing a water-octanol partition coefficient for the CG-ligand and by comparing it with available references. The logarithm of the water-octanol partition coefficient can be computed from the desolvation free energy of the compound in water and octanol as:

$$\log P_{\text{ow}} = \frac{\Delta G_{\text{oco}\rightarrow\text{vac}} - \Delta G_{\text{wat}\rightarrow\text{vac}}}{\ln(10)RT} \tag{1}$$

where R is the ideal gas constant and T is the temperature of the simulated system (300K). Desolvation free energies from water to vacuum $\Delta G_{wat\rightarrow vac}$ and from octanol to vacuum $\Delta G_{oco\rightarrow vac}$ have been computed with `gmx bar` by performing thermodynamic integration over ten simulation windows of 60 ns (of which the first 10 ns have been discarded as equilibration). The error over the computational logP has been estimated starting from the uncertainty over each desolvation free energy $\epsilon_{\Delta G}$ as:

$$\epsilon_{\log \text{P}_{\text{ow}}} = \frac{\sqrt{\epsilon^2_{\Delta G_{\text{oco}\rightarrow\text{vac}}} + \epsilon^2_{\Delta G_{\text{wat}\rightarrow\text{vac}}}}}{\ln(10)RT} \tag{2}$$

Bead type attribution followed forcefield guidelines established in literature[6, 7] and prioritized consistency with previously reported molecules[8]. The $logP_{ow}$ estimated for the CG-model of the ligands are reported in Figure S7b.

**a)**

| Compound | Method | logP | Reference |
|---|---|---|---|
| Combretastatin-A4 | *ALOGPS* | 3.32 | Drugbank |
| | *Chemaxon* | 3.38 | Drugbank |
| | *ACD/logP* | 3.57 | RSC ChemSpider |
| | *log-Kow* | 3.38 | EPI Suite-v4.11 |
| Podophyllotoxin | *ALOGPS* | 2.37 | Drugbank |
| | *Chemaxon* | 1.62 | Drugbank |
| | *ACD/logP* | 1.6 | RSC ChemSpider |
| | *log-Kow* | 2.01 | EPI Suite-v4.11 |
| Guanosine | *Experimental* | -1.9 | Sangster (1993) |
| | *ALOGPS* | -2.1 | Drugbank |
| | *Chemaxon* | -2.7 | Drugbank |
| | *ACD/logP* | -1.7 | RSC ChemSpider |

**b)**

| Compound | $\Delta G_{oco\rightarrow vac}$ [kJ·mol$^{-1}$] | $\Delta G_{wat\rightarrow vac}$ [kJ·mol$^{-1}$] | CG-logP$_{ow}$ |
|---|---|---|---|
| Combretastatin-A4 | 37.14±0.03 | 13.08±0.07 | 4.19±0.01 |
| Podophyllotoxin | 51.75±0.07 | 23.25±0.11 | 4.97±0.02 |
| Guanosine | 45.94±0.12 | 61.29±0.12 | -2.68±0.03 |

**Supplementary Figure S7** a) Reference values for the water-octanol partition coefficient from experimental and chemoinformatics methods. b) Computational $logP_{ow}$ for the proposed Martini 3 parameterization of the compounds simulated.

We note on passing that for combretastatin-A4 we obtain good agreement with chemoinformatics reference, while for podophyllotoxin standard Martini 3 types yield a more lipophilic behavior compared to available references. For GTP, the parameterization has followed two steps. Since the retrieval of reliable partition coefficients for charged moieties is challenging, we opted to first validate the bead types for the neutral guanosine scaffold, obtaining a difference of less than 0.8 logP units with respect to experimental and computational references. Then, for the GTP model, the type for the bead containing the 5' carbon has been modified and standard parameters have been attributed to the charged phosphate groups.

#### 1.2 Protein Modelling

Atomistic parameters of the CHARMM-36m forcefield for the tubulin dimer have been generated with the CHARMM-GUI platform[9, 10] using the crystallographic structure obtained in complex with colchicine (PDB ID 4O2B[11]). After energy minimization with the steepest descent algorithm, a 1 ns NVT equilibration with position restraints on the $C_\alpha$ has been performed (V-rescale thermostat[3], $\tau_T$=1.0 ps), followed by a 10 ns equilibration with the Berendsen barostat[4] ($\tau_P$=5.0 ps, compressibility $4.5 \cdot 10^{-5}\ bar^{-1}$). Finally, a production simulation of 300 ns with the Parrinello-Rahman pressure coupling[5] has been generated. The protocol has been repeated four times with randomization of initial velocities. The stability of the protein's backbone has been verified through the computation of RMSD and RMSF with GROMACS (Figure S8a,c,d). A smooth function Q was used to monitor changes in the fraction of native contacts retained[12] (a metric often used to quantify conformational changes in protein[13])

$$Q = \frac{1}{N_{pairs}} \sum_{p=1}^{N_{pairs}} \frac{1}{1 + exp(\beta(r_P - \lambda r_{p,0}))} \tag{3}$$

Where $N_{pairs}$ is the list of native contacts, $r_P$ the pair distance at time $t$, $r_{p,0}$ the native pair distance, $\beta$ a smoothening factor ( set to 50 $nm^{-1}$[12]) and $\lambda$ a calibration factor (1.8 for AA structures[12], 1.5 for CG structures[14])(Figure S8b).

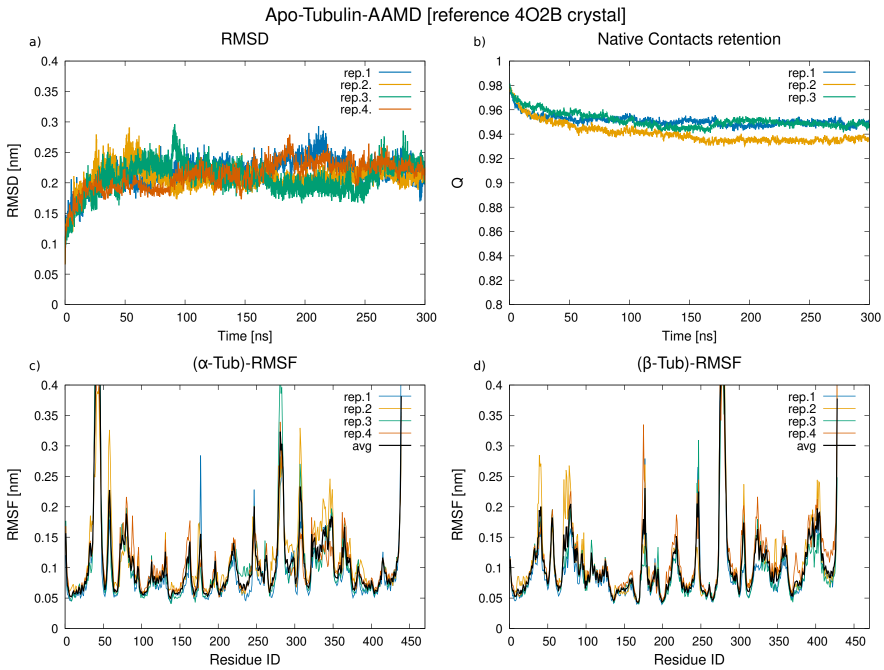

**Supplementary Figure S8** AA-MD assessment of backbone stability for apotubulin starting from the 4O2B crystal structure. a) Backbone RMSD. b) Fraction of crystal contacts retained. Bottom row represents RMSF results for c) $\alpha$-Tubulin and d) $\beta$-Tubulin.

The atomistic results have been taken as a reference to validate the Martini 3 models of the protein. The martinize2 utility[15] was used to generate the CG structure file. For the EN, an upper bond length of 0.85 nm was imposed, and the elastic force constant was set to 750 $kJ\ mol^{-1}$. For Olives and goMartini, default settings have been initially considered. After generation of the Martini 3 topology and insertion of the nucleotides in the GTP E and N sites (see "Nucleotide modelling" in main text for details), the tubulin dimer was solvated with regular water beads and NaCl has been included to neutralize the system's charge and provide a physiological concentration of 0.15 M. After minimization, an equilibration with the Berendsen barostat ($\tau_P$=4.0 ps, compressibility $3 \cdot 10^{-4}\ bar^{-1}$) and position restraints on the backbone beads was performed for 50 ns at 20 fs. Then, three production simulations of 2 $\mu$s have been collected for each protein model (EN, Olives, goMartini) with the Parrinello-Rahman barostat ($\tau_P$=12.0 ps, compressibility $3 \cdot 10^{-4}\ bar^{-1}$). Stability of the CG models for tubulin was assessed by calculating the RMSD and RMSF and by comparing it with atomistic references (Figures S9,S10).

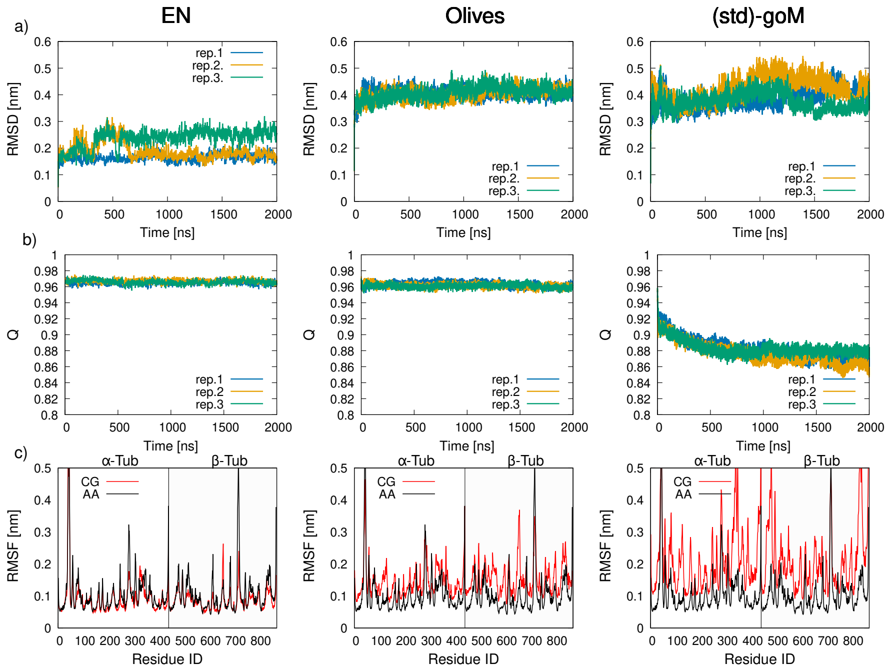

**Supplementary Figure S9** Validation of backbone stability of the CG-models for tubulin. a) RMSD compared to the crystal structure (PDB ID 4O2B) mapped to CG-resolution. b) Retention of the native contacts identified in the crystal structure. c) RMSF analysis: average fluctuations from reference AA-MD simulations are reported in black, while average fluctuations from three CG-MD simulations are reported in red.

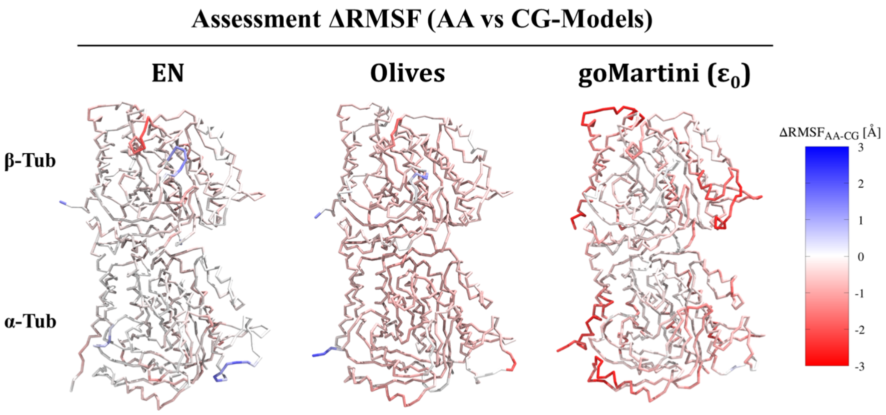

**Supplementary Figure S10** Assessment of $\Delta RMSF$ between reference AA-MD data and CG-models for tubulin. Residues coloured in red fluctuate more in CG compared to AA ($RMSF_{AA}$-$RMSF_{CG} < 0$). Residues in white present negligible differences. Residues in blue are overstabilized by the CG-network compared to AA-MD ($RMSF_{AA} > RMSF_{CG}$).

The goMartini model generated with default settings showed significant residue fluctuations, prompting us to optimize it. A custom protocol has been implemented to redefine the Lennard Jones terms of goMartini based on the reference AA simulations, as summarized by the pseudoalgorithm reported in Algorithm 1.

**Algorithm 1** Reparameterization of GoMartini Lennard–Jones bond terms

**Require:** AA trajectories $\{\mathrm{AA}_i\}$, CG trajectories $\{\mathrm{CG}_j\}$, `go_nbparams.itp`
**Ensure:** Updated CG bond reference distances

$\mathcal{B} \leftarrow$ retrieve_bond_list(`go_nbparams.itp`)
$r_{\mathrm{CG}}^{\mathrm{ref}} \leftarrow$ retrieve_bond_reference(`go_nbparams.itp`)
▷ Map AA trajectories to CG resolution
**for all** $\mathrm{AA}_i \in \{\mathrm{AA}_i\}$ **do**
    $\mathrm{AA}_i^{\mathrm{mapped}} \leftarrow$ map_aa_to_cg($\mathrm{AA}_i$)
**end for**
▷ Compute average CG bond distances
**for all** $b \in \mathcal{B}$ **do**
    **for all** $\mathrm{CG}_j \in \{\mathrm{CG}_j\}$ **do**
        $d_j^{\mathrm{CG}}(b) \leftarrow$ compute_distance($b$, $\mathrm{CG}_j$)
    **end for**
    $\bar{d}^{\mathrm{CG}}(b) \leftarrow \mathrm{average}_j \left[d_j^{\mathrm{CG}}(b)\right]$
**end for**
▷ Compare AA-mapped and CG distances
**for all** $b \in \mathcal{B}$ **do**
    **for all** $\mathrm{AA}_i^{\mathrm{mapped}}$ **do**
        $d_i^{\mathrm{AA}}(b) \leftarrow$ compute_distance($b$, $\mathrm{AA}_i^{\mathrm{mapped}}$)
    **end for**
    $\bar{d}^{\mathrm{AA}}(b) \leftarrow \mathrm{average}_i \left[d_i^{\mathrm{AA}}(b)\right]$
    $\Delta_{\mathrm{ref}} \leftarrow \bar{d}^{\mathrm{AA}}(b) - r_{\mathrm{CG}}^{\mathrm{ref}}(b)$
    **if** $|\Delta_{\mathrm{ref}}| > 0.1$ nm **then**
        update_ref($b$, $\bar{d}^{\mathrm{AA}}(b)$)
    **else**
        $\Delta_{\mathrm{sim}} \leftarrow \bar{d}^{\mathrm{AA}}(b) - \bar{d}^{\mathrm{CG}}(b)$
        **if** $|\Delta_{\mathrm{sim}}| > 0.1$ nm **then**
            update_ref($b$, $\bar{d}^{\mathrm{AA}}(b)$)
        **end if**
    **end if**
**end for**

In particular, the list of the residues involved in the network of LJ-bonds generated by the standard goMartini and their reference distance $d_{CG}^{ref}$ was retrieved from the goMartini itp file. Available AA-simulations of tubulin have been mapped to CG-resolution and for the residue pairs of interest, the average distance $\bar{d}_{AA}$ has been computed. The same average distance has been determined from the CG-simulations with the standard implementation of goMartini ($\bar{d}_{CG}$). Then, for every bond of the goMartini network, a comparison between the target atomistic average ($\bar{d}_{AA}$) and the CG-data ($d_{CG}^{ref}$, $\bar{d}_{CG}$) was made, redefining the $\sigma$ parameter of the LJ-bond to reflect $\bar{d}_{AA}$ if the difference was found to be greater than 0.1 nm. Finally, an increase in the overall bond strength (through the parameter $\epsilon$) of 25% and 50% was considered. This procedure allowed to reduce the overall RMSD with respect to the reference structure and improve the RMSF profile, as depicted in Figure S11. The reparametrized goMartini model with $\epsilon = \epsilon_0 + 50\%$ was found to be particularly stable and to reproduce remarkably well the atomistic fluctuations (Figure S12) and therefore was applied in subsequent simulations.

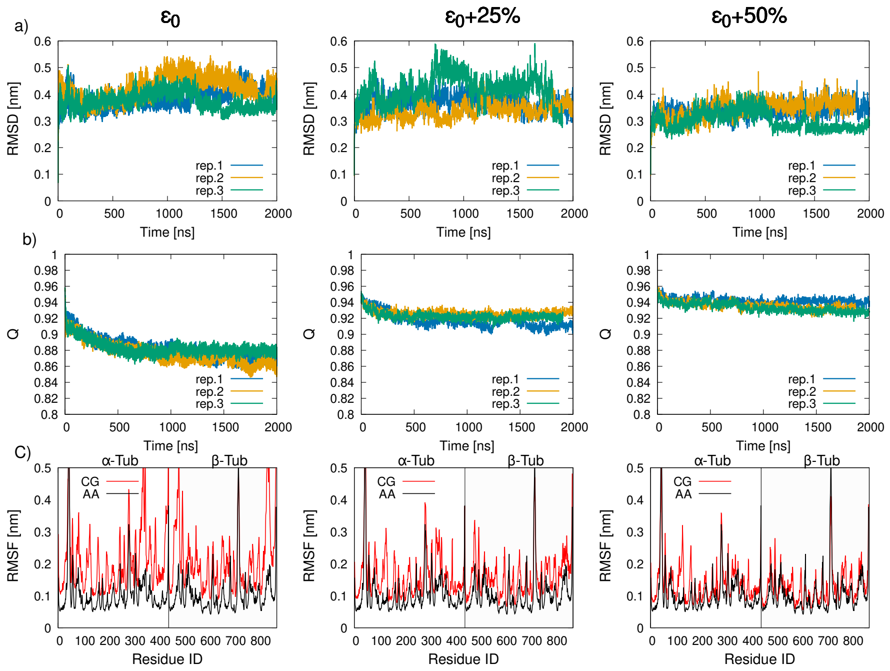

**Supplementary Figure S11** Optimization of the goMartini CG-protein model for the tubulin heterodimer. a) RMSD compared to the crystal structure (PDB ID 4O2B) mapped to CG-resolution. b) Retention of the native contacts identified in the crystal structure. c) RMSF analysis: average fluctuations from reference AA-MD simulations are reported in black, while average fluctuations from three CG-MD simulations are reported in red.

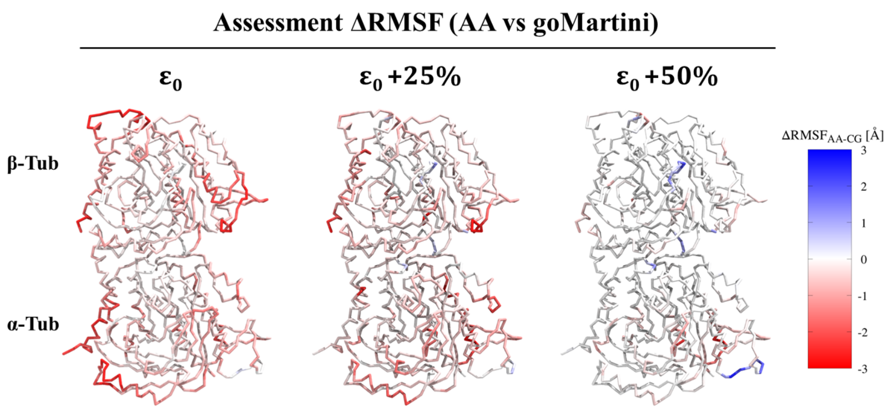

**Supplementary Figure S12** Assessment of $\Delta RMSF$ between reference AA-MD data and the reparameterized goMartini for tubulin. Residues coloured in red fluctuate more in CG compared to AA ($RMSF_{AA}$-$RMSF_{CG}$¡0). Residues in white present negligible differences. Residues in blue are overstabilized by the CG-network compared to AA-MD ($RMSF_{AA} > RMSF_{CG}$).

#### 1.3 Nucleotide Modelling

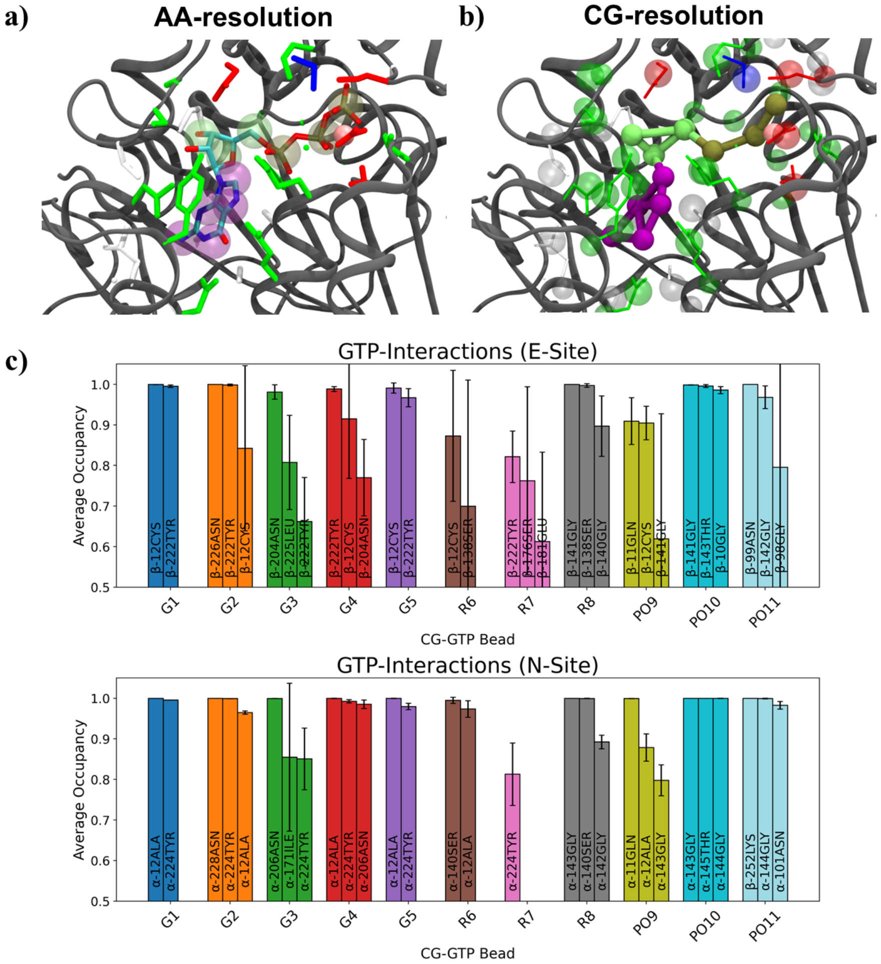

**Supplementary Figure S13** Modelling of GTP-sites in tubulin. a) Atomistic representation of the N-site. Residue sidechains close to GTP are represented in licorice with a color scheme corresponding to their polarity (basic in blue, acidic in red, polar in green and nonpolar in white). Transparent spheres indicate position of the corresponding CG-beads for the Martini 3 model of GTP. B) conversion of the atomistic structure to CG-resolution. GTP has been represented with a ball & stick model, whereas for the nearby interacting protein residues both the atomistic structure (narrow sticks) and the CG (transparent spheres) mapped representation are shown. C) results for the Tubulin-GTP contact analysis performed on the atomistic trajectory mapped to CG-resolution. For each GTP CG-bead, the top three residues interacting most often are reported (if the average occupancy is above 60% of the simulation). Error bar is the standard deviation over three replicas.

#### 1.4 Characterization of ligand binding modes

The characterization of the ligand binding modes starts by performing a contact analysis between the small molecule and the protein using gmx mindist and the standard cutoff of 0.6 nm. The molecule is annotated as bound whenever the number of contacts is greater than 0. If the ligand is annotated as bound for more than 100 ns, the corresponding trajectory file is extracted.
Since a ligand might continuously be annotated as bound when exploring the surface of the protein (Figure S14a), a clustering procedure is applied to identify whether a portion of the trajectory describes a highly localized ligand pose (Figure S14b). A smoothening procedure is applied to merge pose conformationally similar but separated in time by few ns to account for small fluctuations. If the identified subevent lasts still longer than 100 ns, a trajectory is extracted and passed for subsequent analysis.

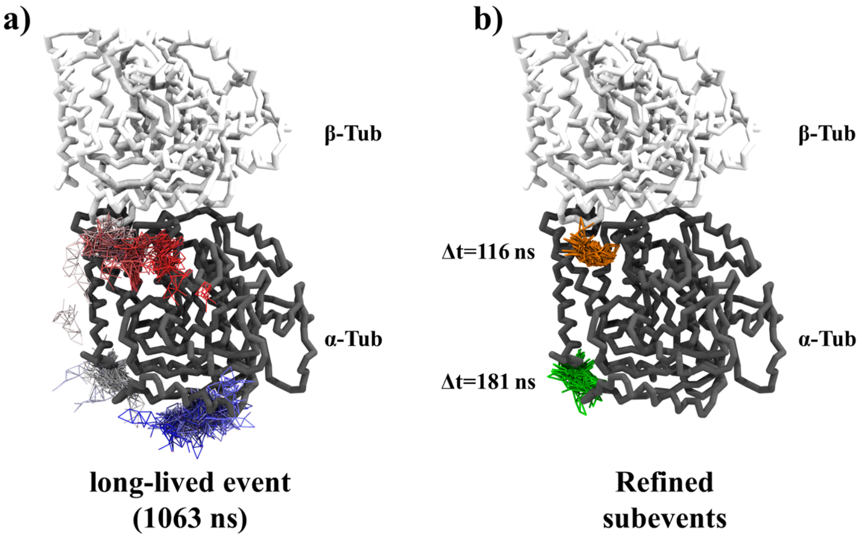

**Supplementary Figure S14** a) Example of continuous long-lived event of CG-colchicine (lasting 1063 ns), exploring the surface of $\alpha$-tubulin. b) Depiction of two highly localized subevents, lasting more than 100 ns each.

In order to identify a conformation representative of the subevent, the gromos algorithm[16] with a cutoff of 0.4 nm is applied. To assess whether the largest cluster can be assumed to be representative of the entire subevent trajectory, the relative population covered by it is considered. If it is below 80%, the subevent is discarded, as it implies that more than one pose contributes significantly. We note on passing that the "Single-linkage" algorithm assigns to the same cluster structure that are distant less than 0.2 nm (in RMSD term). This implies that a series of conformations slowly varying and drifting over time tend to be merged in the same subevent. As a results, the centroid of such collection is not truly representative of the entire population, as is evidenced by Figure S15. Consequently, we applied the gromos algorithm both for the initial identification of the subevents and for the extraction of the centroid representing the subevent. After this processing step, a list of centroid structures files highly representative of an equal number of corresponding distinct trajectories is obtained.

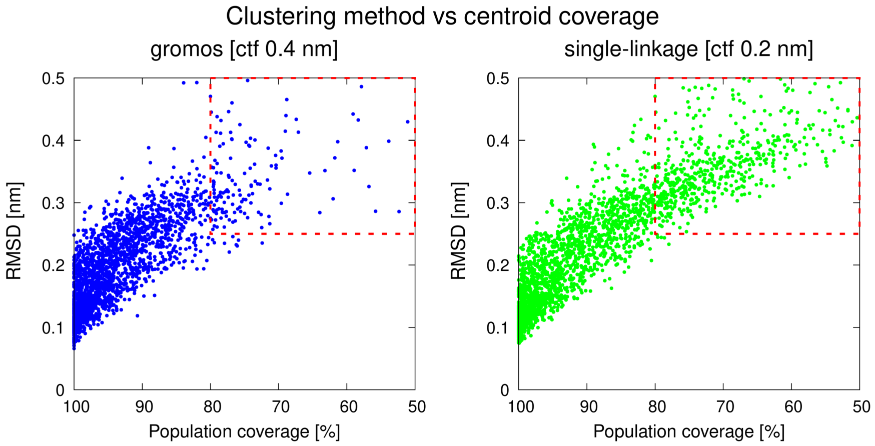

**Supplementary Figure S15** Influence of clustering method used for identifying distinct subevents on the representativeness of the respective centroid. Dashed rectangle identifies a region containing centroids that account for a suboptimal (less than 80%) fraction of their subevent. The "single-linkage" algorithm (reported on the right), yields a high fraction of these poorly representative centroids, whereas the gromos algorithm obtained better results.

The next step is to make a comparison of the poses described by these subevents in order to identify representative binding modes. First a clustering of the centroids with a large cutoff is performed to identify "binding domains". This allows in effect to limit the comparison to ligand poses that are located in the same pocket. Having defined a list of unclassified binding events located in the same region of space, an iterative procedure is applied: the centroid of the first event is taken as reference "supra-cluster" and for every other item of the list, the average RMSD from it is computed. If the average RMSD from the candidate trajectory is below 4 Å, the event is annotated as belonging to the supra-cluster, otherwise it is rejected. At the end of the first iteration, the list is updated to remove all the items attributed and the procedure is repeated, taking the first element of the new list of unclassified items as the reference for the second supra-cluster. At the end of the procedure, for each binding pocket, a list of representative ligand structures with a record of associated binding events is obtained.

Having performed the conformational analysis on the ligand poses, the protocol attempts to characterize the kinetics associated with each Binding Mode (BM). In particular, if more than ten binding events have been collected for a certain pose, an histogram of the event duration will be generated. Many criteria can be followed when building histograms. Currently, the following rules have been implemented

- Freedman-Diaconis's rule (FD)
- Sturger's rule
- Doane's rule

The python function `scypy.optimize.curve_fit` from the library SciPy[17] is then called to fit the probability distribution of the histogram of the binding event durations to the exponential decay $y = Ae^{-\frac{t}{\tau}}$, yielding an estimate for the residence time $\tau$ with an associated uncertainty $\epsilon_\tau$ from the fitting error. For every rule used to generate a histogram, a $\tau$ fit is attempted, allowing for backup in case a particular histogram resulted suboptimal. If less than ten binding events are observed, no estimation attempt is made.

**a)**

| Label | #Events | #Rule | $\tau$ | $\varepsilon_\tau$ |
|---|---|---|---|---|
| D2S1 | 179 | FD | 594.6 | 58 |
| D2S3 | 88 | FD | 1359.7 | 103.5 |
| D7S1 | 15 | FD | 7254.9 | 2672.3 |
| D1S2 | 37 | FD | 1274.1 | 256 |
| D1S1 | 328 | FD | 57 | 3.2 |
| D3S3 | 100 | FD | 236.6 | 20.9 |
| D5S1 | 11 | Doane | 4986.2 | 1908.2 |
| D1S8 | 31 | FD | 1260.8 | 194.6 |
| D2S6 | 74 | FD | 205.9 | 30 |
| D14S4 | 14 | Doane | 3724.4 | 1014.7 |

**b)**

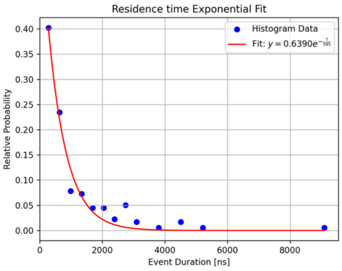

**Supplementary Figure S16** a) Summary of the residence time results for the top-ten BMs identified for the colchicine/(EN)-tubulin biosystem. Second column indicates how many distinct events are associated to the BM. Third column indicates which rule has been applied to generate the histogram used in the fitting procedure. b) Example of residence time estimation by fitting a histogram of the binding events duration.

Finally, for every binding domain a list of relevant ligand BMs with associated residence time is obtained. The results are sorted in order of relative coverage $w_i$ of each binding mode, where the relative coverage is the ratio between the sum of the durations of events $\Delta t_j$ associated to the $BM_i$ and the total duration for all BMs.

$$w_i = \frac{\sum_{j \in B_i} \Delta t_j}{\sum_{B_i} \sum_{j \in B_i} \Delta t_j} \tag{4}$$

A pseudoalgorithm describing the protocol's steps is reported in Algorithm 2.

**Algorithm 2** Identification of binding modes and residence time estimation

**Require:** Set of trajectories $\{T_k\}$, ligand list $\{L_i\}$

**Ensure:** Binding domains, binding modes, and residence times

```
                                ▷ Step 1: Identify long-lived binding events
for all T_k ∈ {T_k} do
    perform contact_analysis(T_k)
    for all detected events in T_k do
        if event.duration > 100 ns then
            T^LL ← extract_event(T_k, start, end)
            add T^LL to refinement list
        end if
    end for
end for
                                ▷ Step 2: Sub-event refinement and smoothing
for all L_i ∈ {L_i} do
    list_subevents_raw ← refine_event(L_i, T^LL)
    list_subevents_smooth ← smoothen(list_subevents_raw)
end for
                                ▷ Step 3: Representative structure selection
for all subevent s ∈ list_subevents_smooth do
    if s.duration > 100 ns then
        centroid ← extract_centroid(s)
        p_rel ← compute_rel_population(centroid, s)
        if p_rel > 0.8 then
            label s as accepted
            add centroid to list_all_centroids
        end if
    end if
end for
                      ▷ Step 4: Identify topographically distinct binding domains
list_binding_domains ← cluster(list_all_centroids)
                ▷ Step 5: Identify representative binding modes within each domain
for all domain d ∈ list_binding_domains do
    unassigned ← centroids in d
    while unassigned not empty do
        supracluster ← select seed from unassigned
        for all candidate ∈ unassigned do
            R ← compute_avg_RMSD(candidate, supracluster)
            if R < 0.4 nm then
                assign candidate to supracluster
                remove candidate from unassigned
            end if
        end for
        add supracluster to list_supracluster[d]
    end while
end for
                                ▷ Step 6: Residence time estimation
for all domain d ∈ list_binding_domains do
    for all supracluster c ∈ list_supracluster[d] do
        if c.nevents > 10 then
            H ← build_histogram(c.event_durations)
            c.residence_time ← fit_restime(H)
        end if
    end for
end for
```

#### 2 Equilibrium CG-MD

**a)**

| Compound | **Colchicine** | **Podophyllotoxin** | **Combretastatin-A4** |
|---|---|---|---|
| αβ-Tubulin | 1 | 1 | 1 |
| Ligand | 6 | 4 | 4 |
| GTP | 2 | 2 | 2 |
| $Mg^{2+}$ | 2 | 2 | 2 |
| $Na^{+}$ | 320 | 320 | 320 |
| $Cl^{-}$ | 288 | 288 | 288 |
| Water-beads | 26522 | 26522 | 26522 |
| Box size | *14.9 nm* | *14.9 nm* | *14.9 nm* |
| Tot beads | 29235 | 29219 | 29207 |

**b)**

| Simulated ligand | Replica duration | **(eq)CG-MD Sampling** replicas / *simulated time* | | |
|---|---|---|---|---|
| | | **EN** | **Olives** | **goMartini** |
| Colchicine | 20.05 μs | 30 / *600 μs* | 30 / *600 μs* | 30 / *600 μs* |
| Podophyllotoxin | 20.05 μs | 35 / *700 μs* | 35 / *700 μs* | 35 / *700 μs* |
| Combretastatin-A4 | 20.05 μs | 35 / *700 μs* | 35 / *700 μs* | 35 / *700 μs* |

**Supplementary Figure S17** Equilibrium CG-MD simulation campaign. a) Size and composition of the simulation boxes. b) Number of replicas collected for each protein-ligand system and total simulated time.

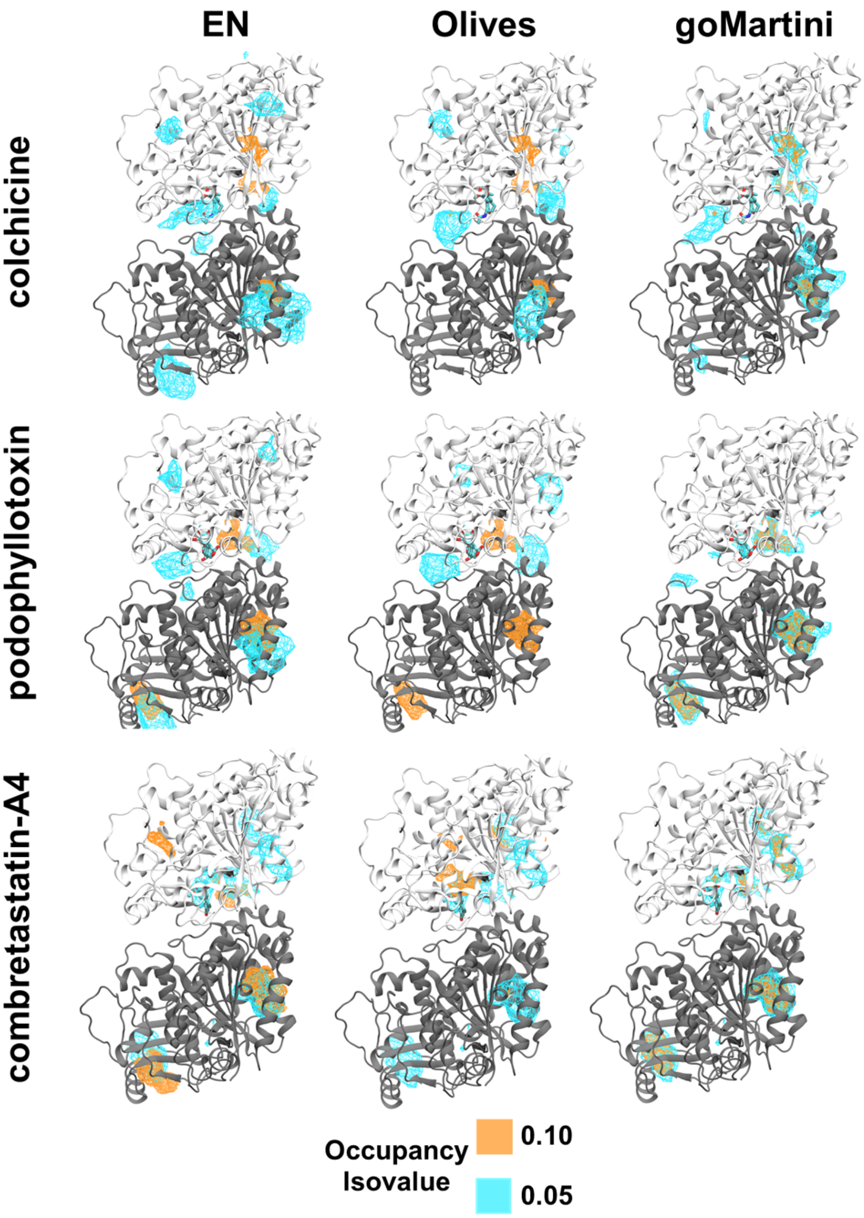

**Supplementary Figure S18** Occupancy isocurves computed from unbiased CG-MD simulations. Each row summarizes the results for each ligand, while every column indicates the CG-protein model considered. $\alpha$-tubulin is colored in gray, while $\beta$-tubulin is depicted in white. Atomistic crystallographic pose is depicted in licorice.

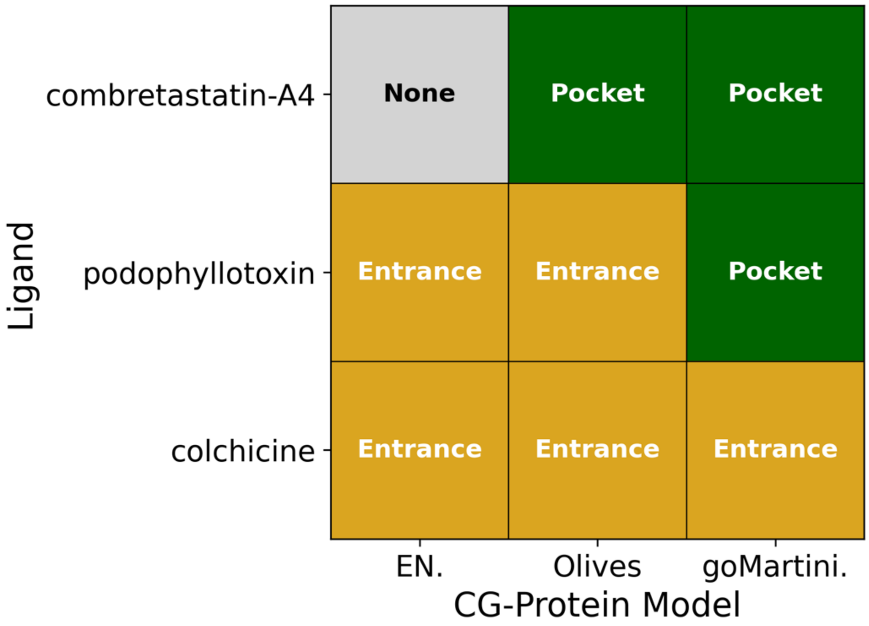

**Supplementary Figure S19** Summary of the relation between top-ten observed poses from unbiased simulation and exploration of the cryptic colchicinoid site. Green indicates that some poses were inside the crystallographic site. Orange indicates that poses were observed at the entrance of the binding channel. Gray indicates that no pose consistent with the experimental information was found.

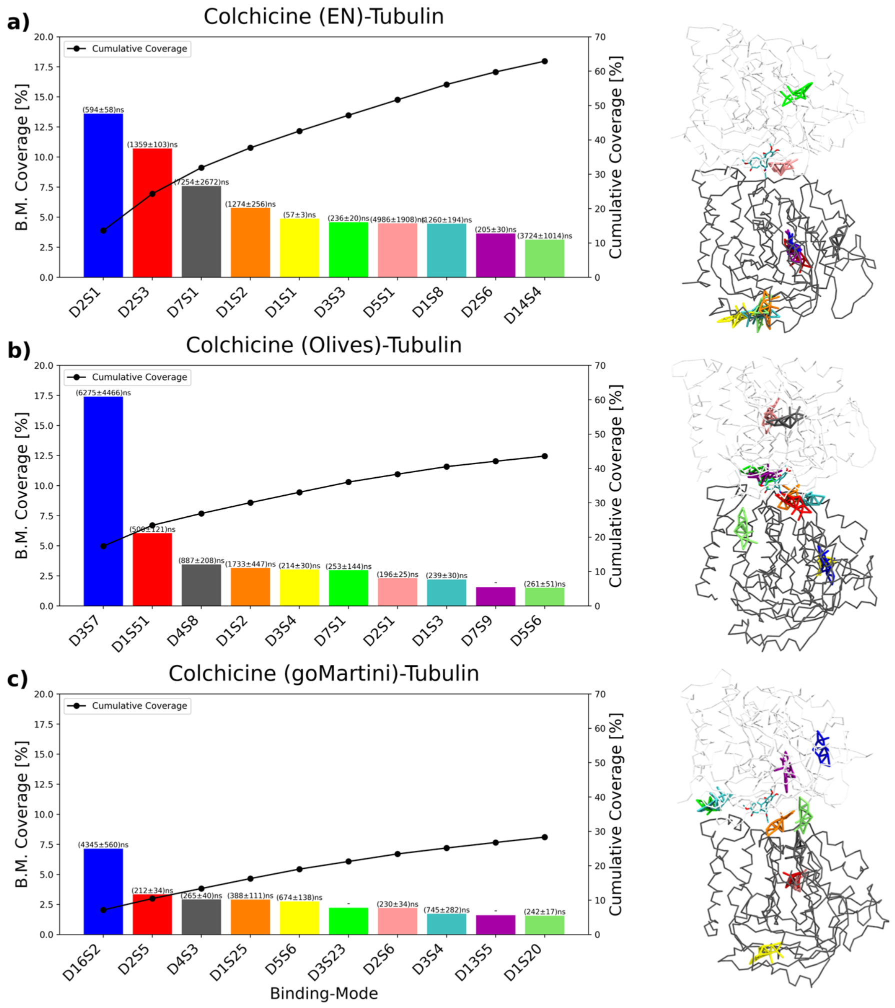

**Supplementary Figure S20** Colchicine binding mode analysis. On the left, quantitative characterization of the ranking and estimated CG-residence time for the different binding modes. On the right, graphical representation of the ranked poses using the same color coding. a) Results for simulations with (EN)-tubulin. b) Results for simulations with (Olives)-tubulin. c) Results for simulations with (goMartini)-tubulin.

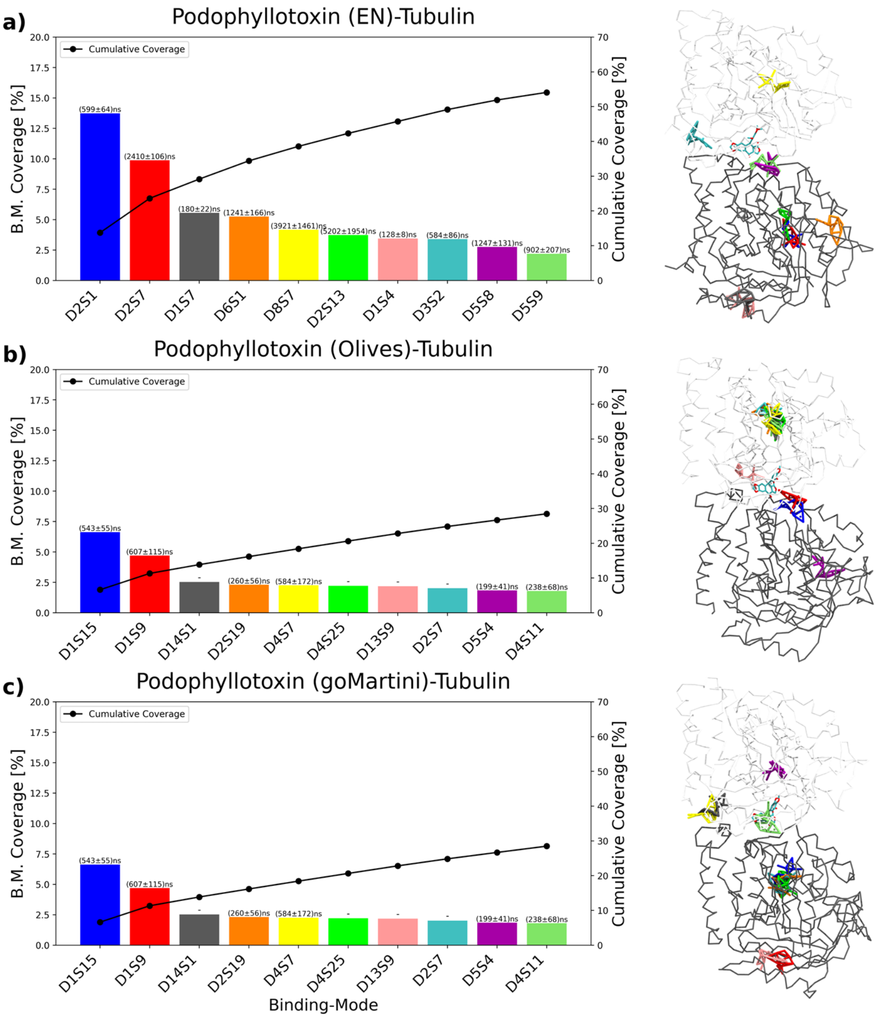

**Supplementary Figure S21** Podophyllotoxin binding mode analysis. On the left, quantitative characterization of the ranking and estimated CG-residence time for the different binding modes. On the right, graphical representation of the ranked poses using the same color coding. a) Results for simulations with (EN)-tubulin. b) Results for simulations with (Olives)-tubulin. c) Results for simulations with (goMartini)-tubulin.

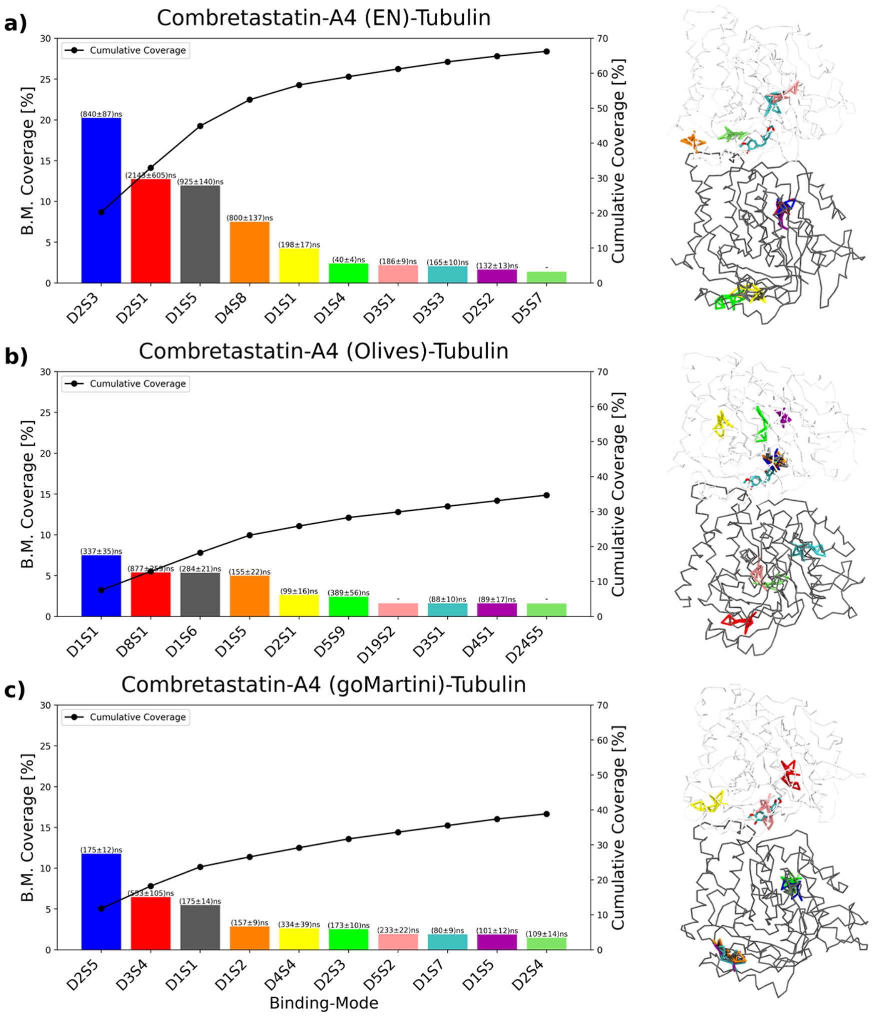

**Supplementary Figure S22** Combretastatin-A4 binding mode analysis. On the left, quantitative characterization of the ranking and estimated CG-residence time for the different binding modes. On the right, graphical representation of the ranked poses using the same color coding. a) Results for simulations with (EN)-tubulin. b) Results for simulations with (Olives)-tubulin. c) Results for simulations with (goMartini)-tubulin.

### 3 Funnel Medadynamics

#### 3.1 FMD Simulation setup

The choice of collective variables (CVs) for the metadynamics setup was designed to maintain transferability to other cases of study. In particular, we identified a set of simple geometric descriptors defining the position of the ligand with respect to the binding pocket, namely the distance between the ligand's center of mass (COM) and the COM of a set of residues lining the interior of the crystallographic site, and the angle among the COM of the residues $\beta$-Leu255, $\beta$-Ala316 and of the ligand (Figure S23).

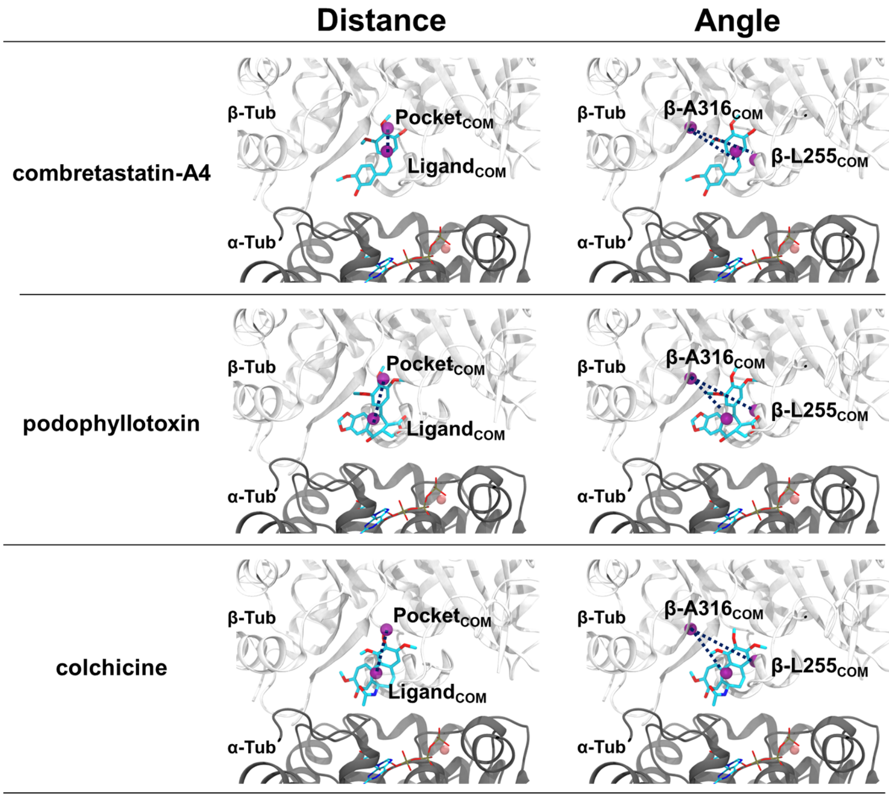

**Supplementary Figure S23** Collective variable definition. Top row: combretastatin-A4; center row: podophyllotoxin; bottom row: colchicine. Left column represents the distance between the ligand and the pocket while the column on the right displays the angle definition.

Initial simulations with AA-FMD proved difficult to converge. In particular, the Free Energy Surface obtained from each simulation showed a topographically distinct location of the energy minima (Figure S24).

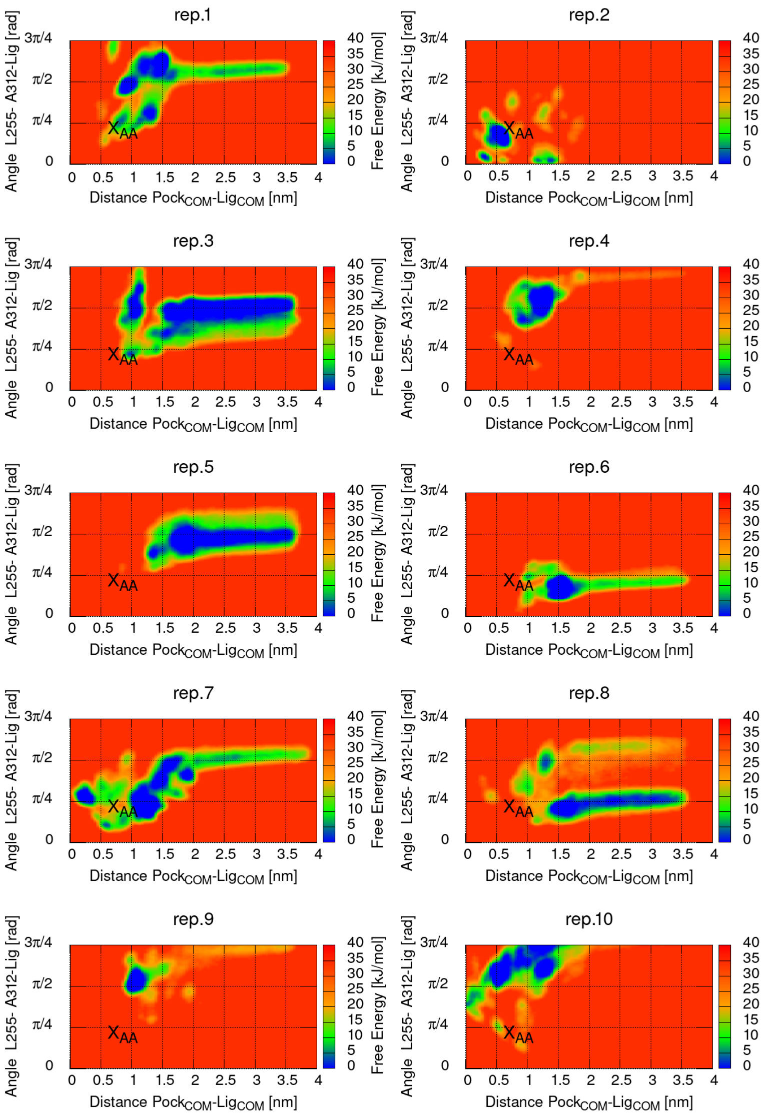

**Supplementary Figure S24** FES obtained from AA-FMD of the tubulin-podophyllotoxin. The denaturation of the loop involved in the CV definition leads to non-converging projection in the CV-space.

Analysis of the trajectory highlighted that under the strain of the metadynamics bias, one of the loops containing a residue involved in the definition of the angular CV ($\beta$-Leu255) underwent denaturation, as highlighted in Figure S25, leading to an ill-definition of the CV and inconsistent FES projection. The introduction of local restraining in the loops containing residues involved in the CVs definition (as detailed in Methods) improved convergence of the results.

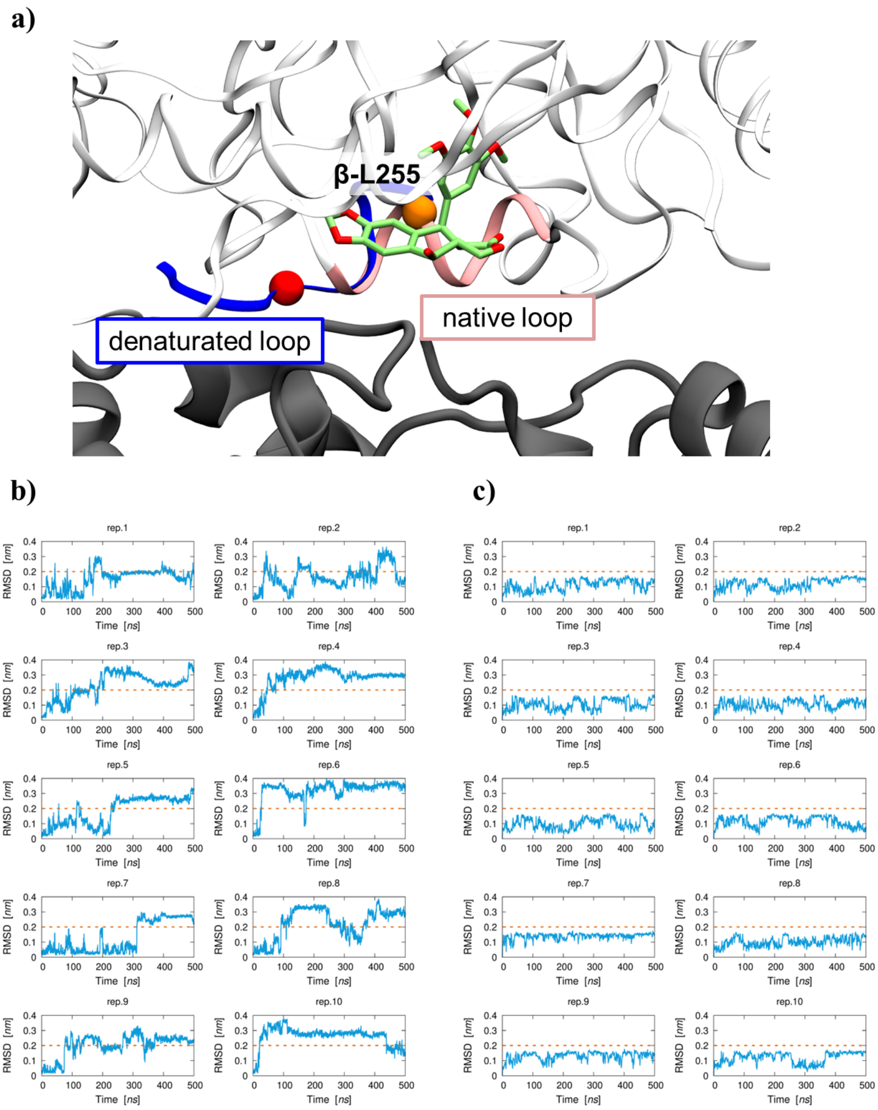

**Supplementary Figure S25** Effects of metadynamics strain on the loops used for the CV definition. a) Denaturation of the loop containing the residue $\beta$-Leu255 ($C_\alpha$ highlighted depicted as a sphere) used to build the angular CV from the native pose (pink) to a deformed conformation (blue). b) RMSD of the loop during ten replicas of AA-FMD without protocol optimization. c) RMSD of the loop after imposition of RMSD upper restraint with PLUMED.

**a)**

| Ligand | AA |
| --- | --- |
| Colchicine | [0.6:1.0] nm / [0.4:0.9] rad |
| Podophyllotoxin | [0.6:1.0] nm / [0.5:0.8] rad |
| Combretastatin-A4 | [0.25:0.8] nm / [0.5:0.8] rad |

**b)**

| Ligand | EN | Olives | goMartini |
| --- | --- | --- | --- |
| Colchicine | [0.4:0.6] nm / [0.5:0.8] rad | [0.4:0.7] nm / [0.5:0.9] rad | [0.5:0.7] nm / [0.4:0.8] rad |
| Podophyllotoxin | [0.4:0.6] nm / [0.5:0.8] rad | [0.5:0.7] nm / [0.5:0.8] rad | [0.6:0.9] nm / [0.4:0.8] rad |
| Combretastatin-A4 | [0.35:0.7] nm / [0.5:1.0] rad | [0.4:0.8] nm / [0.4:0.9] rad | [0.4:0.8] nm / [0.1:0.8] rad |

**Supplementary Figure S26** Basin boundaries for the calculation of free energy from FMD FES. a) CV-intervals for the analysis of AA-FMD results. b) CV-ranges for the analysis of CG-FMD results.

#### 3.2 FMD Simulation campaign

| Ligand | Protein | metadynamics `biasfactor` | Sampling |
|---|---|---|---|
| colchicine | AA-Tubulin | 24 | 10×700 ns |
| colchicine | EN-Tubulin | 12 | 18×10 μs |
| colchicine | EN-Tubulin | 24 | 18×10 μs |
| colchicine | EN-Tubulin | 50 | 16×10 μs |
| colchicine | Olives-Tubulin | 12 | 16×10 μs |
| colchicine | Olives-Tubulin | 24 | 12×10 μs |
| colchicine | Olives-Tubulin | 50 | 12×10 μs |
| colchicine | goMartini-Tubulin | 12 | 10×10 μs |
| colchicine | goMartini-Tubulin | 24 | 12×10 μs |
| colchicine | goMartini-Tubulin | 50 | 11×10 μs |
| podophyllotoxin | AA-Tubulin | 24 | 10×500 ns |
| podophyllotoxin | EN-Tubulin | 12 | 12×10 μs |
| podophyllotoxin | EN-Tubulin | 24 | 12×10 μs |
| podophyllotoxin | EN-Tubulin | 50 | 12×10 μs |
| podophyllotoxin | Olives-Tubulin | 12 | 12×10 μs |
| podophyllotoxin | Olives-Tubulin | 24 | 12×10 μs |
| podophyllotoxin | Olives-Tubulin | 50 | 11×10 μs |
| podophyllotoxin | goMartini-Tubulin | 12 | 12×10 μs |
| podophyllotoxin | goMartini-Tubulin | 24 | 12×10 μs |
| podophyllotoxin | goMartini-Tubulin | 50 | 16×10 μs |
| combretastatin-A4 | AA-Tubulin | 24 | 10×500 ns |
| combretastatin-A4 | EN-Tubulin | 12 | 12×10 μs |
| combretastatin-A4 | EN-Tubulin | 24 | 12×10 μs |
| combretastatin-A4 | Olives-Tubulin | 12 | 12×10 μs |
| combretastatin-A4 | Olives-Tubulin | 24 | 11×10 μs |
| combretastatin-A4 | goMartini-Tubulin | 12 | 12×10 μs |
| combretastatin-A4 | goMartini-Tubulin | 24 | 12×10 μs |

**Supplementary Table S1** Funnel Metadynamics simulation campaign. For each ligand, first entry refers to a collection of AA-FMD simulations, while the remainder are CG-FMD simulations with different CG-protein networks.

#### 3.3 FMD Results

##### 3.3.1 Influence of CG-protein model on the binding free energy landscape

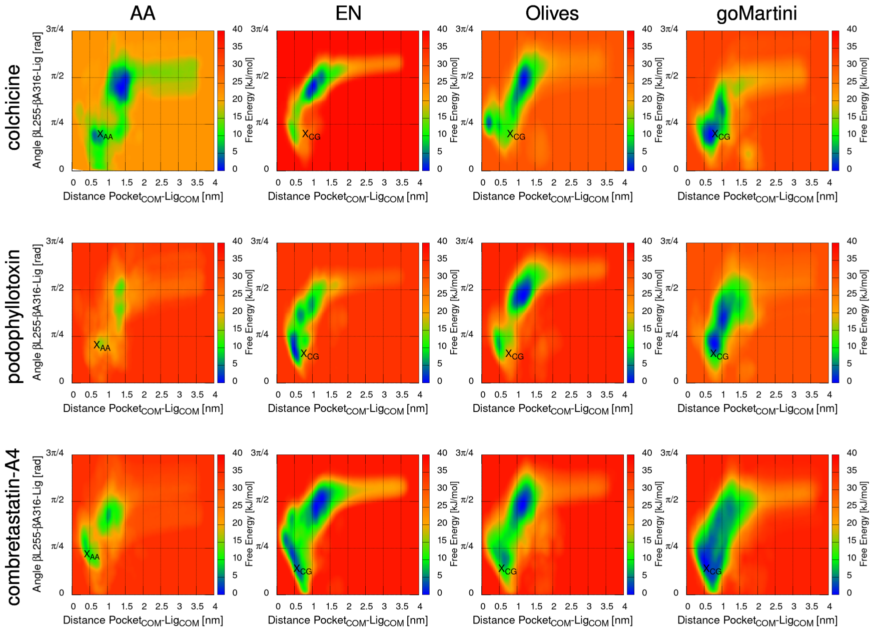

**Supplementary Figure S27** Free Energy Surfaces (FESs) obtained from Funnel Metadynamics. Each row corresponds to a different ligand. First column contains results from AA-FMD. 2nd to 4th columns represent data from CG-FMD performed with different CG-networks applied to the backbone of the tubulin dimer. $X_{AA}$ marks the projection of the crystallographic pose in the CV-space. $X_{CG}$ indicates the projection of the crystal pose mapped to Martini 3 resolution.

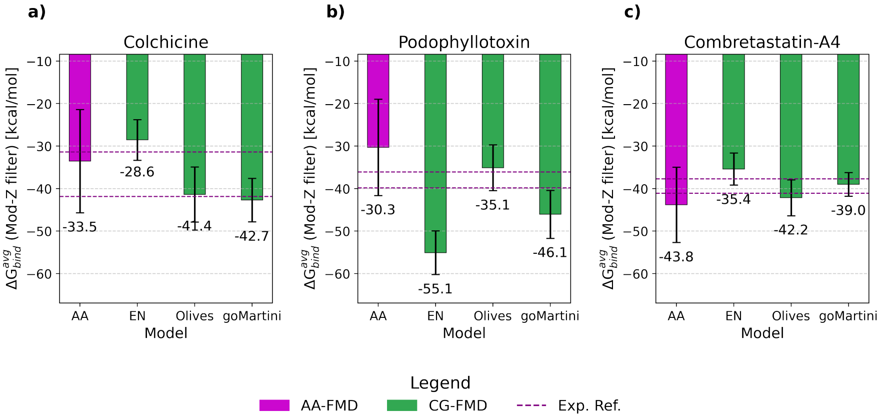

**Supplementary Figure S28** Summary of the binding free energy estimates obtained for the three ligands investigated in this work: a) colchicine, b) podophyllotoxin, c) combretastatin-A4. Value reported in purple was obtained with AA-FMD, while results in green derive from CG-FMD applied to the three CG-protein networks applied to tubulin. Error bars represent the standard error of the mean. Dashed lines indicate the maximum and minimum experimental values reported in literature (summarized in Table S4)

| Ligand | Type | $\Delta G_{\text{bind}}$ [kJ mol$^{-1}$] Half-duration | $\Delta\Delta G_{\text{bind}}^{\text{exp-FMD}}$ [kJ mol$^{-1}$] Half-duration | $\Delta G_{\text{bind}}$ [kJ mol$^{-1}$] Full-duration | $\Delta\Delta G_{\text{bind}}^{\text{exp-FMD}}$ [kJ mol$^{-1}$] Full-duration |
|---|---|---|---|---|---|
| Colchicine | Experimental | [−32:−41] | – | [−32:−41] | – |
| | AA-FMD | $-17 \pm 12$ | −19 | $-33 \pm 12$ | −2.8 |
| | CG-FMD (EN) | $-37 \pm 2$ | 0.8 | $-29 \pm 4$ | −7.7 |
| | CG-FMD (Olives) | $-27 \pm 8$ | −9.2 | $-41 \pm 4$ | 5.1 |
| | CG-FMD (goMartini) | $-54 \pm 6$ | 18 | $-43 \pm 5$ | 6.4 |
| | M.A.E. (CG-FMD) | – | 9.2 | – | 6.4 |
| Podophyllotoxin | Experimental | [−36:−40] | – | [−36:−40] | – |
| | AA-FMD | $-32 \pm 11$ | −5.3 | $-30 \pm 11$ | −6.5 |
| | CG-FMD (EN) | $-53 \pm 12$ | 14.8 | $-55 \pm 5$ | 17.4 |
| | CG-FMD (Olives) | $-41 \pm 10$ | 3.8 | $-35 \pm 5$ | −2.6 |
| | CG-FMD (goMartini) | $-48 \pm 3$ | 10.8 | $-46 \pm 6$ | 8.4 |
| | M.A.E. (CG-FMD) | – | 9.8 | – | 9.5 |
| Combretastatin-A4 | Experimental | [−38:−41] | – | [−38:−41] | – |
| | AA-FMD | $-72 \pm 20$ | 32.8 | $-44 \pm 9$ | 4.2 |
| | CG-FMD (EN) | $-33 \pm 5$ | −6.2 | $-35 \pm 4$ | −4.2 |
| | CG-FMD (Olives) | $-45 \pm 5$ | 5.8 | $-42 \pm 4$ | 2.6 |
| | CG-FMD (goMartini) | $-42 \pm 5$ | 2.2 | $-39 \pm 3$ | −0.6 |
| | M.A.E. (CG-FMD) | – | 4.7 | – | 2.5 |

**Supplementary Table S2** Summary of binding free energy predictions obtained with Funnel Metadynamics (FMD). For each ligand, results obtained both at the AA and CG resolution are reported, alongside the reference experimental value. Uncertainty over the computational results is the standard error of the mean across the available independent replicas. As preliminary indication of convergence, results obtained at half simulation duration and at full length are reported. For CG-results, a mean absolute error (MAE) across the different protein networks is reported.

##### 3.3.2 Optimization of the biasfactor parameter

The solidity of the CG-FMD simulation protocol has been assessed by investigating the influence of metadynamics decrease parameter biasfactor. Reassuringly, we found that the binding free energy predictions converge to similar values (Figure S29), typically within an interval comparable to the statistical uncertainty associated with each average free energy value. We note on passing that in the case of colchicine binding to Olives-tubulin, the application of a soft `biasfactor=12` led to a poor sampling of the binding pocket and a significant underestimation of the free energy, hence the choice to highlight `biasfactor=24` as a more robust setup value which performed more consistently across all simulated biosystem.

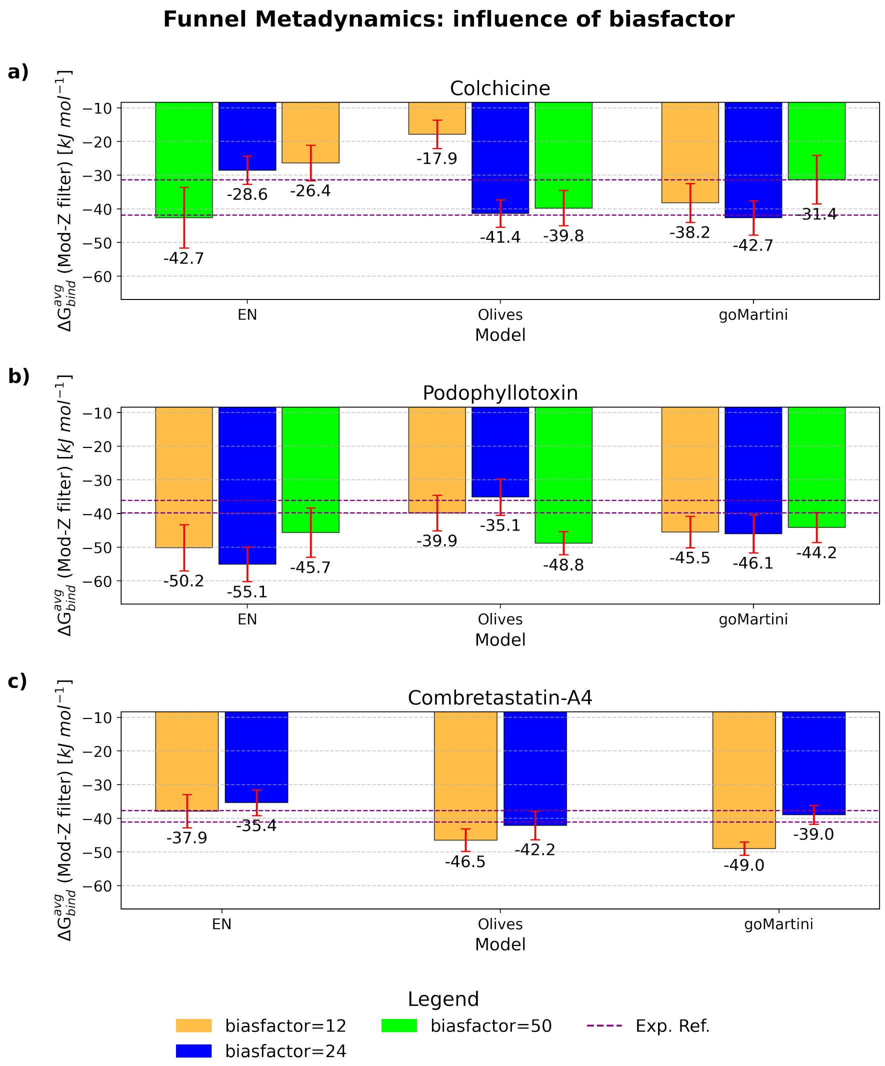

**Supplementary Figure S29** Investigation of the effect of the biasfactor metadynamics parameter on the final estimate of the binding free energy. Panel a) summarizes results for colchicine, b) for podophyllotoxin, c) for combretastatin-A4. Error bar indicates SEM across available replicas.

##### 3.3.3 FMD-Convergence of the simulations

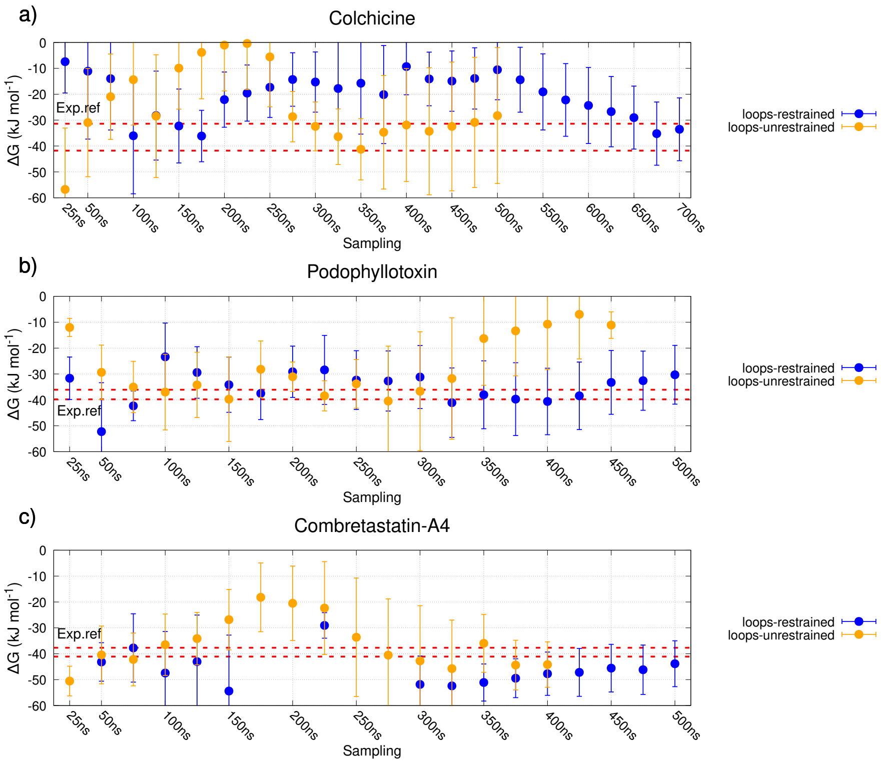

**Supplementary Figure S30** Convergence of binding free energy estimation with AA-FMD. Results are presented in the following order: a) colchicine, b) podophyllotoxin, c) combretastatin-A4. Error bar represents the SEM over the available replicas. Dashed line indicates experimental affinity reference.

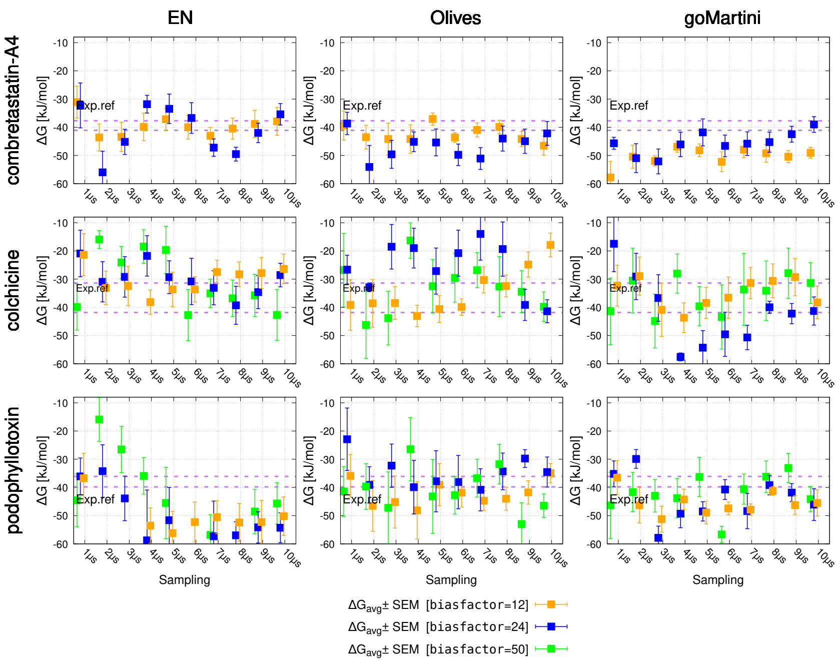

**Supplementary Figure S31** Convergence in time of the binding free energy estimation from CG-FMD. Each row represents a distinct ligand, while each column represents a different protein network applied to tubulin's backbone. In each sublot, results obtained with different biasfactor parameter value are reported. Error bar represents the SEM, across available replicas.

#### 3.4 Reference values for protein ligand binding affinity

| Protein | Ligand | Type | $\Delta G_{\text{bind}}$ [kJ mol$^{-1}$] | $\lvert\Delta\Delta G_{\text{bind}}^{\text{exp}-\text{FMD}}\rvert$ [kJ mol$^{-1}$] | Reference |
|---|---|---|---|---|---|
| BRD4 | Colchicine | Experimental | -27 | - | [18] |
| BRD4 | Colchicine | AA-FMD | -36±4 | 9 | [19] |
| BRD4 | PYCA | Experimental | -28 | - | [18] |
| BRD4 | PYCA | AA-FMD | -35±4 | 7 | [19] |
| BRD4 | SPCA | Experimental | -38 | - | [18] |
| BRD4 | SPCA | AA-FMD | -65±7 | 27 | [19] |
| BRD4 | RVX-OH | Experimental | -37 | - | [20] |
| BRD4 | RVX-OH | AA-FMD | -48±7 | 11 | [19] |
| BRD4 | RVX-208 | Experimental | -38 | - | [20] |
| BRD4 | RVX-208 | AA-FMD | -37±13 | 1 | [19] |
| Colchicalin | Colchicine | Experimental | -57 | - | [21] |
| Colchicalin | Colchicine | AA-FMD | -49±8 | 8 | [19] |
| acetylcholinesterase | metformin | Experimental | -15 | - | [22] |
| acetylcholinesterase | metformin | AA-FMD | -7±1 | 8 | [23] |
| Methionine-AP-II | Ligand-1 | Experimental | -48 | - | [24] |
| Methionine-AP-II | Ligand-1 | AA-FMD | -36 | 12 | [25] |
| Androgen receptor | DHT | Experimental | -51 | - | [26] |
| Androgen receptor | DHT | AA-FMD | -53±0.1 | 2 | [26] |

**Supplementary Table S3** Summary of protein-ligand binding free energy estimations computed with AA-FMD recently reported in literature.

| Ligand | $K_d$ [$\mu$M] | $\Delta G_{\text{bind}}$ [kJ mol$^{-1}$] | Reference |
|---|---|---|---|
| colchicine | 4.92 | $-31.5 \pm 0.4$ | Banerjee *et al.*[27] |
| colchicine | $3.7 \pm 0.18$ | $-32.2$ | Yamazaki *et al.*[28] |
| colchicine | $0.5 \pm 0.9$ | $-37.4$ | Tahir *et al.*[29] |
| colchicine | 0.3 | $-38.6$ | Bhattacharyya *et al.*[30] |
| colchicine | 0.1 | $-42$ | Diaz *et al.*[31] |
| podophyllotoxin | 0.83 | $-36.1$ | Tahir *et al.*[29] |
| podophyllotoxin | 0.5 | $-37.4$ | Kelleher *et al.*[32] |
| podophyllotoxin | $0.2 \pm 0.3$ | $-39.8$ | Wilson *et al.*[33] |
| combretastatin-A4 | $4.00 \pm 0.06$ | $-37.7$ | Lin *et al.*[34] |
| combretastatin-A4 | $1.8 \pm 0.3$ | $-40.0$ | Tahir *et al.*[29] |
| combretastatin-A4 | 1.2 | $-41.1$ | Woods *et al.*[35] |

**Supplementary Table S4** Summary of experimental binding affinity estimates reported in literature for colchicine, podophyllotoxin, combretastatin-A4. Estimated uncertainty is reported only when provided by the original authors

### 4 Equilibrium CG-MD exploration of the pocket

| Simulated ligand | Replica duration | EN | Olives | goMartini |
|---|---|---|---|---|
| colchicine | 20 $\mu$s | 10 | 10 | 10 |
| podophyllotoxin | 10 $\mu$s | 10 | 10 | 10 |
| combretastatin-A4 | 20 $\mu$s | 10 | 10 | 6 |

**Supplementary Table S5** Simulation campaign for the equilibrium CG-MD simulations starting with the ligand already inside the cryptic pocket. For each CG-protein model, the number of collected replicas is reported.

Colchicine eq-CGMD [inside binding site]

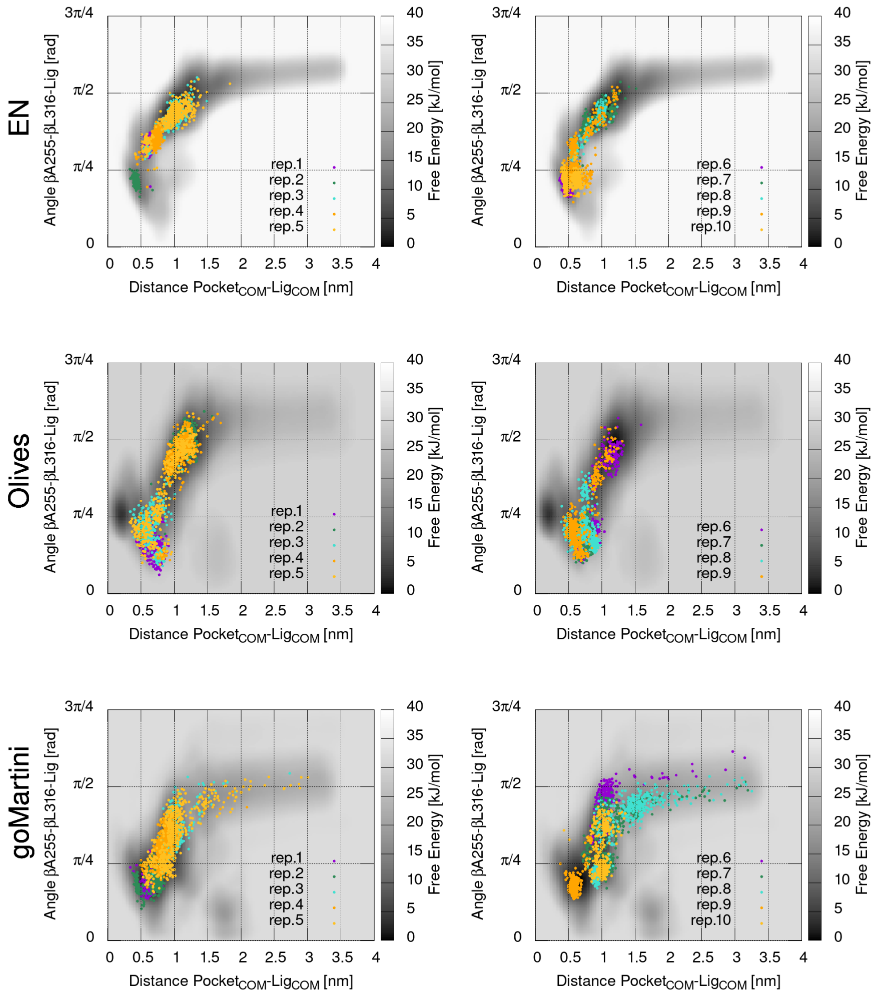

**Supplementary Figure S32** Overlap between the CV projection from equilibrium CG-MD simulation of colchicine starting from inside the binding site (colored data points) and the FES predicted from CG-FMD (grayscale). Each row represents a different CG-protein model applied to the tubulin dimer's backbone. Plots on the left report data from replicas 1-5, while on the right results from replicas 6-10 are presented.

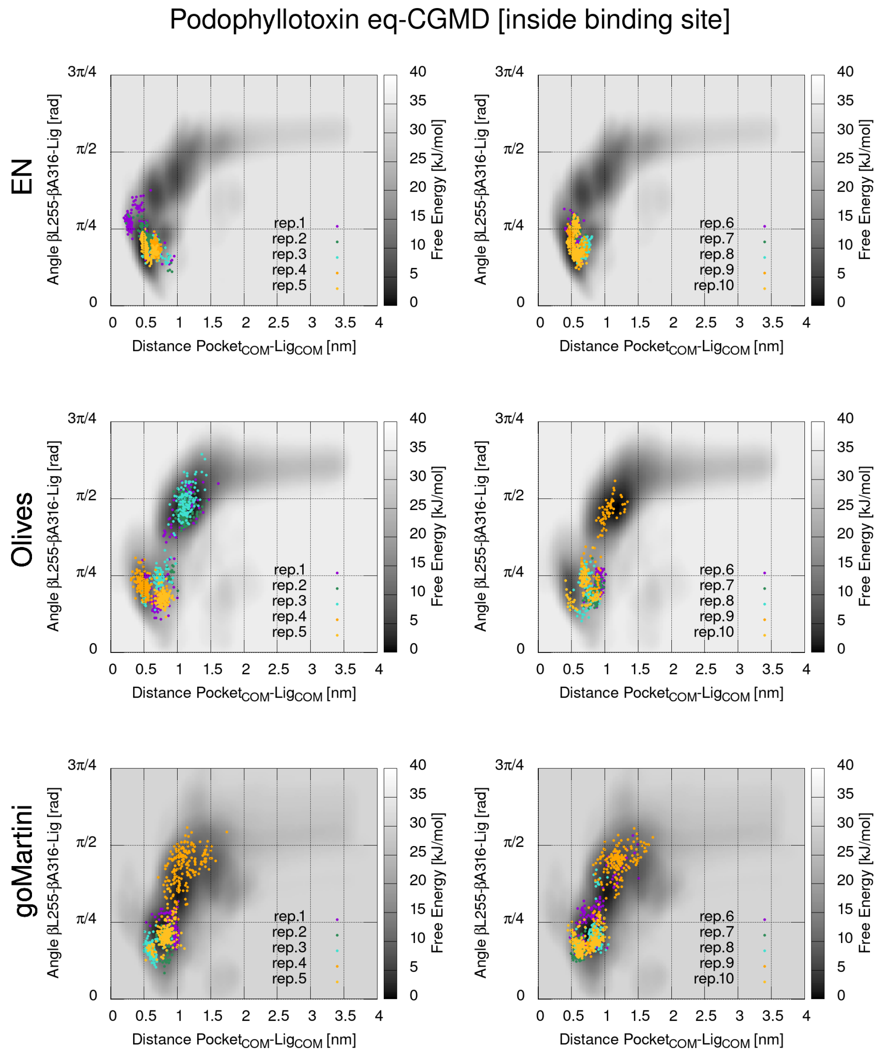

**Supplementary Figure S33** Overlap between the CV projection from equilibrium CG-MD simulation of podophyllotoxin starting from inside the binding site (colored data points) and the FES predicted from CG-FMD (grayscale). Each row represents a different CG-protein model applied to the tubulin dimer's backbone. Plots on the left report data from replicas 1-5, while on the right results from replicas 6-10 are presented.

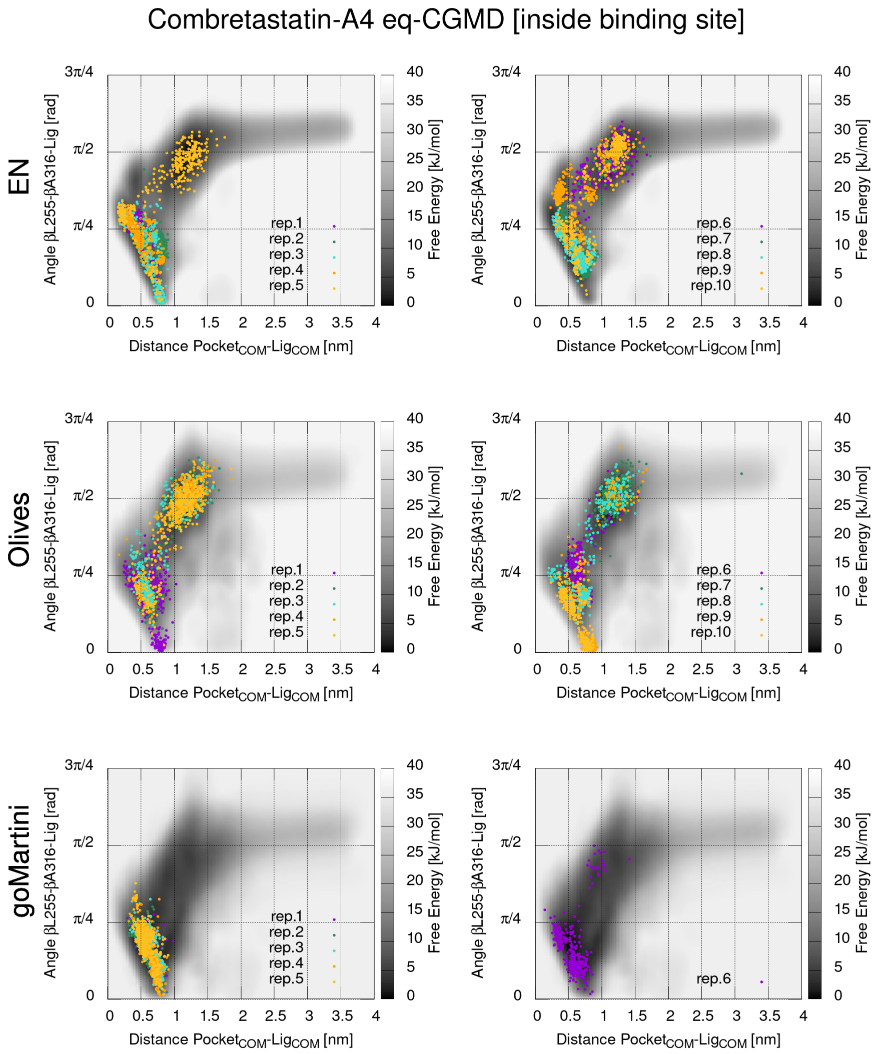

**Supplementary Figure S34** Overlap between the CV projection from equilibrium CG-MD simulation of combretastatin-A4 starting from inside the binding site (colored data points) and the FES predicted from CG-FMD (grayscale). Each row represents a different CG-protein model applied to the tubulin dimer's backbone. Plots on the left report data from replicas 1-5, while on the right results from replicas 6-10 are presented (for goMartini only six replicas were available).

### 5 Assessment of simulation efficiency

a)

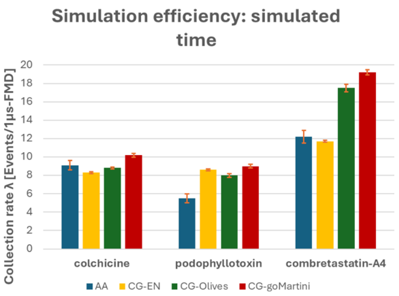

b)

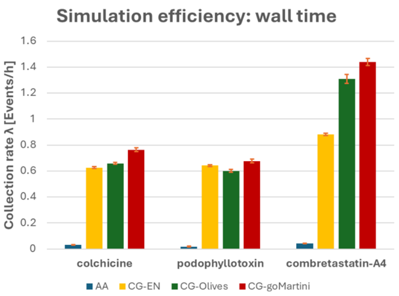

c)

| *Ligand* | AA-FMD | | CG-FMD (EN) | | CG-FMD (Olives) | | CG-FMD (goMartini) | |
|---|---|---|---|---|---|---|---|---|
| | λ [Events/1µs-FMD] | $R^2$ fit | λ [Events/1µs-FMD] | $R^2$ fit | λ [Events/1µs-FMD] | $R^2$ fit | λ [Events/1µs-FMD | $R^2$ fit |
| colchicine | 9.1±0.5 | 0.964 | 8.3±0.1 | 0.998 | 8.8±0.1 | 0.998 | 10.2±0.2 | 0.998 |
| podophyllotoxin | 5.5±0.5 | 0.938 | 8.6±0.1 | 0.998 | 8.0±0.2 | 0.992 | 9.0±0.2 | 0.992 |
| combretastatin-A4 | 12.2±0.7 | 0.970 | 11.7±0.1 | 0.998 | 17.5±0.4 | 0.988 | 19.2±0.3 | 0.988 |

d)

| *Ligand* | AA-FMD | | CG-FMD (EN) | | CG-FMD (Olives) | | CG-FMD (goMartini) | |
|---|---|---|---|---|---|---|---|---|
| | λ [Events/h] | $R^2$ fit | λ [Events/h] | $R^2$ fit | λ [Events/h] | $R^2$ fit | λ [Events/h] | $R^2$ fit |
| colchicine | 0.030±0.002 | 0.964 | 0.625±0.007 | 0.998 | 0.659±0.007 | 0.998 | 0.764±0.013 | 0.994 |
| podophyllotoxin | 0.018±0.002 | 0.938 | 0.642±0.007 | 0.998 | 0.601±0.012 | 0.992 | 0.677±0.014 | 0.992 |
| combretastatin-A4 | 0.041±0.002 | 0.970 | 0.88±0.009 | 0.998 | 1.31±0.034 | 0.988 | 1.439±0.026 | 0.994 |

**Supplementary Figure S35** Determination of the binding event collection rate by linear regression. The first row presents a comparison of the simulation efficiency between AA-FMD and CG-FMD both in terms of a) virtually simulated time ($t_{md} = n_{md-steps} \cdot dt$) and of b) physical wall time. Results from the fitting procedure ($\lambda \pm std.err$, $R^2$) are reported both in terms of c) virtually simulated time and of d) physical wall time.

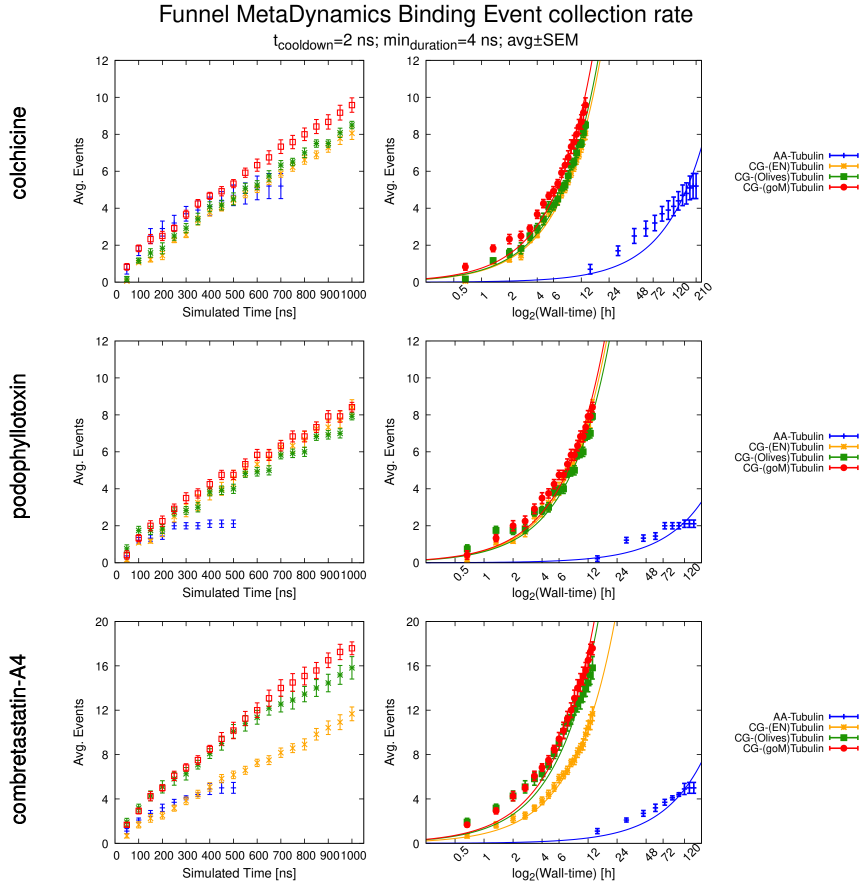

**Supplementary Figure S36** Determination of binding event collection rate by linear regression. Each row represents a distinct tubulin binder. On the left, binding events observed are plotted as a function of the simulated time ($t_{md} = n_{md-steps} \cdot dt$). On the right, the binding events observed are reported as a function of the wall time. Note that a log2 scale is applied, to emphasize that CG-FMD events are collected within few hours, whereas AA-FMD events require days of simulation.
